# Mutant ATXN1 impacts human and mouse microglia and contributes to cognitive, mood, and motor deficits in SCA1 mice

**DOI:** 10.64898/2026.02.10.705104

**Authors:** Adem Selimovic, Gourango Talukdar, Gavin J. Fuchs, Vamika Sharma, Khadija N. Abbas, Sriyan C. Reddy, Eshaan Parnerkar, Ian M. Brooks, Ying Zhang, Michael Koob, Yasushi Nagakawa, Harry Orr, Marija Cvetanovic

## Abstract

Microglia, resident immune cells of the brain, are important players in neurodegeneration. While microglial activation is a hallmark of many neurodegenerative diseases, the specific role of microglia intrinsic factors in microglial activation and disease pathogenesis remains unknown. Spinocerebellar ataxia type-1 (SCA1) is an inherited autosomal dominant neurodegenerative disease characterized by severe neuronal loss and early microglial activation in the cerebellum. SCA1 is caused by CAG repeat expansion in the ubiquitously expressed *ATAXIN1* (*ATXN1)* gene. Using human microglia differentiated from SCA1 patient derived iPSCs, we found that mutant *ATXN1* is sufficient to alter morphology, gene and protein expression in human microglia in a cell-autonomous manner. Moreover, compared to controls, human SCA1 microglia exhibited increased phagocytosis and pro-inflammatory cytokine production, indicating an immune priming. To determine the extent to which mutant *ATXN1* in microglia contributes to SCA1 pathogenesis and behavioral symptoms, we removed mutant *ATXN1* from microglia and macrophages in a novel conditional SCA1 mouse model, *f-ATXN1^146Q/2Q^* mice. Microglial mutant *ATXN1* reduction led to a marked correction in microglia phenotype, in particular in the transcriptomic signature of interferon type 1 mediated immune response, reduced microglial density and resulted in smaller microglia with reduced branching in the cerebellum. Pathology of Purkinje neurons and cerebellar astrogliosis were also ameliorated. Utilizing a battery of behavioral tests, we found that microglia and macrophage mutant *ATXN1* reduction ameliorated cognitive, mood, and motor deficits in SCA1 mice. Together, these results indicate that mutant *ATXN1* directly impacts microglial phenotype in SCA1, contributing to SCA1 pathology and behavioral deficits.

## Introduction

Spinocerebellar ataxia type-1 (SCA1) is an inherited autosomal dominant neurodegenerative disease caused by 39-84 polyglutamine (polyQ) encoding CAG repeats within the *ATAXIN1 (ATXN1)* gene (Klinke et al. 2010; Koeppen 2005; Seidel et al. 2012; Orr 2012; Paulson et al. 2017). SCA1 symptoms include progressive cerebellar ataxia, dysarthria, executive dysfunction and premature death 10-20 years following symptom onset, with longer CAG repeat number resulting in earlier onset and more severe symptoms (Jacobi et al. 2011; Opal and Ashizawa 1998; Ashizawa et al. 2013, 2001; Bürk et al. 2001). SCA1 is characterized by profound loss of Purkinje cells and gliosis in the cerebella of SCA1 patients (Opal and Ashizawa 1998; Ashizawa et al. 2013; Robitaille et al. 1997). While mutant *ATXN1 (mATXN1)* is ubiquitously expressed, most studies focus on *mATXN1* impact on neurons, and how mutant ATXN1 impacts glial cells remains unknown.

Microglia are the resident immune cells that play important roles in brain physiology and pathology by phagocytosing cell debris and synapses, secreting trophic factors and cytokines (Gallo et al. 2022; Nakayama et al. 2018; Favuzzi et al. 2021; Araki et al. 2021; Kreisel et al. 2019; Marín-Teva et al. 2004; Augusto-Oliveira et al. 2019). In neurodegeneration, microglia undergo a process of activation resulting in changes in density, morphology, phagocytosis, cytokine secretion and gene expression. Activated microglia exhibit both neuroprotective and neurotoxic phenotypes and have been characterized by spectrum of gene expression of several different phenotypes including neurotoxic M1, neuroprotective M2 and disease associated microglia (DAM) (Hickman et al. 2018; Liao et al. 2012; Guo et al. 2022; Parhizkar et al. 2019). In SCA1, cerebellar microglia exhibit signs of activation, such as increased density and transcriptional changes from the early stages of disease (Tejwani et al. 2024; Cvetanovic 2015; Qu et al. 2017; Rosa et al. 2023). Removing the majority of microglia with pharmacological inhibition of CSFR1 slightly improved motor deficits in SCA1 mice indicating that microglia may have neurotoxic phenotype in SCA1 (Qu et al. 2017). While *mATXN1* is known to perturb transcription and is expressed in microglia, how *mATXN1* impacts microglia and the extent to which *mATXN1* expression in microglia contributes to their phenotype in SCA1 remains unknown.

In this study, we first investigated how *mATXN1* expression impacts human microglia generated from induced pluripotent stem cells (iPSCs) from SCA1 patients and unaffected sibling controls. SCA1 human microglia exhibited larger soma size, were more phagocytic and secreted more pro-inflammatory cytokines compared to control microglia. This indicated that *mATXN1* expression is sufficient to cause pro-inflammatory microglial phenotype in a cell-autonomous manner. We then investigated the extent to which microglial *mATXN1* contributes to microglial phenotype and disease pathogenesis in SCA1 mice. We used a novel conditional SCA1 model, *f-ATXN1^146Q/2Q^* mice in which coding regions of human ATXN1 with 146 CAG repeats are surrounded by LoxN sites and knocked into one of the endogenous mouse *Atxn1* allele (Duvick et al. 2024). We crossed *f-ATXN1^146Q/2Q^* mice with *Lyve1^CRE^* mice, in which Cre recombinase is highly expressed in microglial precursor cells and macrophages (Pham et al. 2010; Scott et al. 2022), to generate SCA1 mice with selective deletion of *mATXN1* in brain microglia (*ATXN1_mKO_*). *ATXN1_mKO_* had ameliorated gene expression, microglial and cerebellar pathology, motor, cognitive and mood deficits. Our findings highlight the importance of *mATXN1* expression in microglia on SCA1 pathogenesis.

## Materials and Methods

### Generation, characterization, and maintenance of iPSC cells

Skin fibroblasts were obtained from participants after written informed consent, in accordance with protocols approved by the Institutional Review Board of the Human Subjects Committee at the University of Minnesota. Induced pluripotent stem (iPS) cells were generated from fibroblasts (Tolar et al. 2011) using the CytoTune-iPS Sendai Reprogramming 2.0 Kit (Invitrogen), following the manufacturer’s instructions. iPS cell characterization was performed as previously described (Tolar et al. 2011). The length of CAG repeats in the ATXN1 gene was analyzed for each expanded allele to confirm pathogenic expansion (Sheeler et al. 2024). The used hiPSC’s lines were in table-1:

**Table-1.**

| ID | Sex | Disease | WT repeat | Expanded repeat |
| --- | --- | --- | --- | --- |
| 11-033 | M | SCA1 | 32 | 43 |
| 11-034 | M | unaffected | nd |  |
| 11-036 | M | SCA1 | 30 | 41 |
| 11-037 | M | SCA1 | 33 | 46 |
| 11-038 | M | unaffected | 29 |  |
| 11-039 | F | unaffected | nd |  |

Human iPSCs were cultured in mTeSR™ plus medium (STEMCELL^TM^ Technologies, Cat. No. 100-0276) with 1% of pen/strep (Gibco, Cat, No. 15140-148) on growth factor–reduced Matrigel® (Corning, Cat. No. 354230)–coated culture vessels. Cells were maintained at 37 °C with 5% CO₂ in a humidified incubator. The culture medium was replaced daily using pre-warmed mTeSR™ plus medium. Cells were passaged every 5–7 days at a split ratio of approximately 1:6, following manufacturer-recommended dissociation and reseeding procedures. iPSC cultures used for experiments were passaged 2 to 3 times after thawing.

### Differentiation and maturation of microglia from hiPSCs

Human induced pluripotent stem cells (hiPSCs) were maintained under feeder-free conditions in mTeSR™ plus medium Matrigel®–coated culture vessels until reaching approximately 70–80 % confluence. Differentiation toward the microglial lineage was performed using STEMCELL Technologies reagents in a two-stage process that follows the manufacturer’s documentation for the STEMdiff™ Hematopoietic Kit (Cat. No. 05310) and STEMdiff™ Microglia Differentiation and Maturation Kits (Cat. Nos. 100-0019 and 100-0020).

In the first stage, hiPSC colonies were exposed to the STEMdiff™ Hematopoietic differentiation medium to generate suspension cultures enriched in hematopoietic progenitor cells (HPCs). The appearance of non-adherent, round progenitors was monitored visually as described in the kit guide. At the manufacturer-recommended time point (Day-12), HPCs were collected and resuspended in the basal medium supplied with the STEMdiff™ Microglia Differentiation Kit to initiate commitment to the microglial precursor lineage (Douvaras et al. 2017).

During the second stage, these precursors were transferred to low-attachment plates containing the microglia differentiation medium. After the appearance of small, motile, ramified cells consistent with microglial morphology, cultures were transitioned to the STEMdiff™ Microglia Maturation Medium (Cat. No. 100-0020) to promote functional maturation on Day-24. Cultures were maintained in a humidified incubator at 37 °C with 5 % CO₂, with medium exchanges performed according to the manufacturer’s protocol. Maturation continued until cultures exhibited stable microglia-like morphology and marker expression (Douvaras et al. 2017). For characterization, differentiated cells were fixed and analyzed by immunocytochemistry or flow cytometry for canonical microglial markers on Day-30 to Day-34. All procedures were conducted in accordance with institutional biosafety and ethical regulations. Detailed procedural steps, reagent compositions, and timing were outlined in the STEMCELL Technologies product manuals referenced above.

### Flow cytometry

For flow cytometric analysis, hiPSC-derived HPCs and microglia were gently dissociated into a single-cell suspension using an enzyme-free cell dissociation solution, as recommended in the STEMdiff™ Microglia differentiation Kit documentation (STEMCELL Technologies). Cells were washed and resuspended in phosphate-buffered saline (PBS) containing 1% bovine serum albumin (BSA). For surface antigen detection, cells were incubated with fluorochrome-conjugated primary antibodies against HPC and microglial markers, including CD43 (Biolegend, Cat. No. 343206), CD45 (Biolegend, Cat. No. 982305), CD11b (Biolegend, Cat. No. 982608), and pHrodo (green) bioparticles (Thermo Fisher Scientific, Cat. No. P35366) for phagocytosis. For intracellular staining, the receptor of the cells was blocked with Human TruStain FcXTM (Biolegend, Cat. No. 422301) according to the manufacturer’s recommendations, followed by incubation with antibodies recognizing HPCs and microglia. Stained cells were analyzed on a BD LSRFortessa™ or equivalent flow cytometer, and data were processed using FlowJo™ software (BD Biosciences). Fluorescence compensation and gating were performed using single-stained controls and fluorescence-minus-one (FMO) controls. At least 10,000 viable single-cell events for phagocytosis and 20,000 viable single-cell events for HPCs and microglia were recorded per sample for analysis.

### Immunocytochemistry (ICC)

Cells for fluorescent immunolabeling were grown on glass coverslips. Plated cells were fixed using freshly prepared 10% formalin in PBS for 20 minutes at RT. Following fixation, cells were washed 1X with RT PBS before permeabilization with 0.25% PBS-Triton for 10 minutes. Blocking was performed in 10% NGS for 1hr at RT. Primary antibody against microglial marker ionized calcium binding adapting molecule 1 (Iba1) (Abcam, Cat. No. AB107159) in 4% NGS were applied overnight at 4°C followed by 3X wash in PBS at RT. Finally, secondary antibodies (Anti Rabbit-Alexa 594) were applied for 1 hr at RT in a light-free environment before a final 3X wash in PBS and rinse in diH^2^O. Coverslips were airdried in a light-free environment and plated on glass slides using Vectashield with DAPI (Cat. No. H-1500).

### pHrodo™ Green Phagocytosis Assay

Phagocytic activity was assessed using pHrodo™ Green–labeled bioparticles according to the manufacturer’s instructions (Invitrogen, Cat. No. P35366). The pHrodo™ dye exhibits minimal fluorescence at neutral pH and fluoresces brightly in the acidic environment of phagosomes and lysosomes, enabling selective detection of internalized particles. Bioparticles were added to microglial cultures and incubated under standard culture conditions to allow phagocytosis, which was quantified by measuring green fluorescence using flow cytometry and fluorescence microscopy.

### Human cytokine array

A human cytokine array was performed using the Proteome Profiler™ Cytokine Array Kit (Bio-Techne, R&D Systems, USA; Cat. No. ARY005B). Cell culture supernatants (two control and two SCA1 samples, with two replicates each) were collected and centrifuged at 400 × g for 5 minutes at 4 °C to remove debris. The clarified supernatants were aliquoted and stored at −80 °C until analysis. Prior to use, samples were thawed on ice and diluted with an array buffer according to the manufacturer’s instructions.

The cytokine array was performed following the manufacturer’s protocol. Briefly, nitrocellulose membranes were blocked with Array Buffer 1 for 1 hr at room temperature on a rocking platform. Prepared samples were mixed with the biotinylated detection antibody cocktail and incubated for 1 hour at room temperature. The sample–antibody mixtures were then added to the membranes and incubated overnight at 4 °C with gentle agitation.

Following incubation, membranes were washed three times with a wash buffer to remove unbound proteins. Membranes were then incubated with streptavidin–horseradish peroxidase (HRP) for 2.5 hrs at RT on a rocking platform shaker. After additional washing steps, cytokine signals were detected using the provided chemiluminescent substrate, Super Signal Femto (Thermo Fisher Scientific), and visualized and analyzed using an ImageQuant LAS 4000.

### LC–MS/MS-based microglial proteomic analysis

Cell pellets were lysed in proteomic lysis buffer (7 M urea, 2 M thiourea, 0.4 M Tris pH 8, 20% acetonitrile, 10 mM TCEP, 40 mM chloroacetamide), sonicated, incubated at 37 °C, and clarified by centrifugation. Protein concentrations were determined by Bradford assay, and 1 µg of protein per sample was digested overnight with trypsin (1:40, w/w) at 37 °C. Peptides were acidified with formic acid, desalted using SDB-RPS StageTips (Rappsilber et al. 2003), and dried by vacuum centrifugation.

Dried peptides were reconstituted in 0.1% formic acid and analyzed (100 ng per sample) by capillary LC–MS/MS with a Thermo Fisher Scientific, Inc (Waltham, MA) Dionex UltiMate 3000 RSLCnano system coupled to an Orbitrap Eclipse mass spectrometer (Thermo Scientific, Waltham MA). Peptides were separated on a self-packed C18 column using a linear acetonitrile gradient and analyzed in data-independent acquisition (DIA) mode with HCD fragmentation. MS1 spectra were acquired in the Orbitrap at 60,000 resolution, and MS2 spectra at 30,000 resolutions.

Peptide tandem MS data were processed using CHIMERYS 3.0.0 (https://www.msaid.de/) within Proteome Discoverer 3.1 against the human UniProt database (UP000005640, August 2025) supplemented with a common contaminant database (https://github.com/HaoGroup-ProtContLib).

Search was performed with trypsin specificity, up to two missed cleavages, carbamidomethylation of cysteine as a fixed modification, and methionine oxidation as a variable modification. Label-free quantification was performed in Proteome Discoverer using MS2-based fragment ion intensities of unique and razor peptides, with normalization to summed peptide abundances. Protein ratios were calculated using a MaxLFQ-like approach (Cox et al. 2014), and statistical significance was assessed using a background t-test with Benjamini–Hochberg false discovery rate correction. 1% protein and peptide False Discovery Rate (FDR) and 2 minimum peptides were applied as filters on the protein report using the Percolator algorithm (Käll et al. 2007) in PD.

### Mice

The creation of *f-ATXN1^146Q/2Q^* mice was previously described (Duvick et al. 2024). These knock-in mice express coding regions of human mutant *ATXN1* surrounded by LoxN sites, under the endogenous mouse promoter, allowing for cell specific removal of mutant *ATXN1* via Cre expression.

The creation of *Lyve1^CRE^* mice was previously described (Pham et al. 2010). These knock-in mice express EGFP-hCre under the *Lyve1* promoter. In all experiments investigators were blinded to the genotype of the mice. Animal experimentation was approved by the Institutional Animal Care and Use Committee (IACUC) of University of Minnesota and was conducted in accordance with the National Institutes of Health’s (NIH) Principles of Laboratory Animal Care (86–23, revised 1985).

### RNA-sequencing

Total RNA was extracted from human microglia using a PureLink RNA mini kit (Invitrogen, Cat. No. 12183018A) according to the manufacturer’s instructions. Briefly, microglial cells were lysed using the provided lysis buffer to ensure complete disruption of cellular membranes and inactivation of RNases. For mice, RNA was extracted from flash frozen cerebellums using TRIzol Reagent (Life Technologies). RNA was sent to the University of Minnesota Genomics Center for quality control, including fluorometric quantification (RiboGreen assay, Life Technologies) and RNA integrity with capillary electrophoresis (Agilent BioAnalyzer 2100, Agilent Technologies Inc.). All submitted samples with RNA integrity numbers (RINs) 7.1 or greater proceeded to RNA sequencing on an TruSeq Stranded Total RNA using 150 nt paired-end read strategy (Hamel et al. 2024). Data is stored on University of Minnesota Supercomputing Institute Servers. To analyze raw data, paired short reads were analyzed through CHURP (Collection of Hierarchical UMII-RIS Pipelines, PEARC ‘19 Proceeding, Article No. 96), which includes data quality control via FastQC, data preprocessing via Trimmomatic (Bolger et al. 2014), mapping via HiSat2 (Dagher et al. 2015) against reference mouse genome and expression quantification via Subread (Liao et al. 2014). The resulting count matrix of gene expression was used as input of R (https://www.r-project.org/) package: DESeq2 (v1.44.0) (Love et al. 2014), which tested the differential gene expression. Genes with resulting P values <0.05 were considered significant. DESeq2 variance stabilizing transformation was used to determine outliers. Pathway genes were determined using gProfiler (https://biit.cs.ut.ee/gprofiler/gost). Pathway analysis and dot plot creation was performed using the clusterProfiler package (v4.12.6) using the top 10 pathways in Gene Ontology Biological Processes and Transcription Factors based on P values <0.05 and LogFC <-0.6 and >0.6. Heatmaps were created using the pheatmap package (https://CRAN.R-project.org/package=pheatmap) (v1.0.12).

### Reverse transcription quantitative polymerase chain reaction (RT-qPCR)

Total RNA was extracted following microglia enrichment using PureLink RNA mini kit (ThermoFisher, Cat. No.12183018A) on preseparation, flowthrough, and enrichment phases. RT-qPCR was performed as previously described (Hamel et al. 2024). For cerebellar extracts, we used IDT primetime primers for *Trim5*, *Trim12a*, *Trim30D*. 18S RNA (Forward: AGTCCCTGCCCTTTGTACACA, Reverse: CGATCCGAGGGCCTCACTA) was used for normalization. For validation of enrichment, we used IDT primetime primers for *Aif1* (IDT assay ID: Mm.PT.58.32501510), *Ncam2* (IDT assay ID: Mm.PT.58.9403828), *Slc1a3* (IDT assay ID: Mm.PT.58.6069702) and as above 18S RNA for normalization. For assessment of mutant *ATXN1* knock-out, we used IDT primetime primer human *ATXN1* (IDT assay ID: Hs.PT.58.38425463) and Rpl13a (IDT assay ID: Mm.PT.58.43276970.g) for normalization.

### Microglia Enrichment

Microglia were enriched using Adult Brain Dissociation kit (Miltenyi Biotec, Cat. No. 130-107-677) and following provided protocol. Briefly, littermate-controlled mice 4 to 20 weeks of age were euthanized using CO_2_ and brains were extracted. Brains were homogenized using gentleMACS™ Octo Dissociator with Heaters (Miltenyi Biotec, Cat. No. 130-134-029). Following homogenization solutions underwent debris clearing and red blood cell removal. Finally, microglia were enriched using CD11b MicroBeads (Miltenyi Biotec, Cat. No. 130-093-636) and magnetic LS columns (Miltenyi Biotec, Cat. No. 130-042-401). Cells were separated into three phases for analysis: pre-separation phase (20% of the cells prior to magnetic separation), a flowthrough phase, and an elution phase. Pre-separation phase was used to assess the number of cells obtained from brain dissociation using a hemocytometer.

### Immunofluorescent (IF) staining

We used five littermate and sex-matched mice of each genotype (N = 5 each of *f-ATXN1^146Q/2Q^*, *ATXN1_mKO_*, and wild-type (*WT^2Q/2Q^*), at 34-36 weeks of age. IF was performed on a minimum of three floating 40-μm-thick brain slices from each mouse. Investigators performing staining, imaging and analysis were blinded for the genotype until all quantification was completed. All mice within the cohort were stained at the same time. We used the primary antibody Iba1 (rabbit, Abcam, Cat. No. AB107159), primary antibodies against Purkinje cell marker calbindin (rabbit, Cell Signaling Technology, Cat. No. D1I4Q), vesicular glutamate transporter 2 (VGLUT2) (guinea pig, Millipore, Cat. No. AB2251-I), astrocytic marker glial fibrillary acidic protein (GFAP) (chicken, Millipore, Cat. No. AB5541) as previously described (Hamel et al. 2024).

### Imaging

Confocal images of min of three images per slice per mouse were acquired using a Leica Stellaris 8 microscope using an oil 20x objective. Z-stacks consisting of twenty non-overlapping 1-μm-thick slices were taken of each lobule X per brain slice. The laser power and detector gain were standardized and kept consistent between mice within a cohort, and all images within a cohort were taken in one imaging session. Analysis was conducted using a cell counter in ImageJ (NIH). Thresholding settings were applied and kept consistent across all images prior to automated cell counting.

### Microglia morphology quantification

Three-dimensional reconstructions of microglia were generated using the “filaments” function in Imaris 10.2 (Oxford Instruments). Approximately 1 to 5 microglia were traced from each confocal image from cerebellar lobule X. Microglia that were not wholly in frame were excluded from analysis. Filament parameters were kept consistent for all analysis and intensity parameters were standardized amongst confocal images that came from the same staining batch and microscope session.

### Statistical analysis

Wherever possible, sample sizes were calculated using power analyses based on the standard deviations from our previous studies, significance level of 5%, and power of 90%. Statistical tests were performed with GraphPad Prism 10.0 or JMP Student Edition 18.2.2. Data from immunohistochemistry imaging was analyzed using either a Student’s t-test or one-way ANOVA with Tukey’s post-hoc test depending on whether the comparison was between genotypes or within genotypes. DEGs for RNA sequencing data were identified by a P threshold of <0.05 and Log2FC >0.6 or <-0.6. RNA sequencing pathway analysis was conducted using DEGs Log2FC >0.6 or <-0.6 and P value <0.05. Behavior was analyzed using either one-way or two-way ANOVA with Tukey’s post-hoc test depending on the number of variables compared.

## Results

### Mutant ATXN1 is sufficient to alter morphology and function of human microglia in a cell-autonomous manner

To determine how mATXN1 impacts human microglia, we used patient and unaffected sibling control derived induced pluripotent stem cells (iPSCs) to create SCA1 and control human microglia. We first generated hematopoietic progenitor cells (HPCs) from iPSC lines and used flow cytometry to ensure that HPCs are ≥ 90% CD43+ before proceeding to differentiation (Figure 1A, Supplementary Figure 1A). We then differentiated HPC into microglia using STEMdiff™ Microglia Differentiation and Microglia Maturation Mediums (Figure 1A). Flow cytometry with antibodies against microglial markers CD45 and CD11b demonstrated more than 90% of cells positive for these markers in both SCA1 and control microglia indicating successful differentiation (Supplementary Figure 1B). Using immunofluorescence (IF) with anti-Iba1 antibody, we found that SCA1 human microglia had decreased sphericity and increased soma size compared to unaffected control microglia (Figure 1 C-D), consistent with activated microglia phenotype.

**Figure 1.**
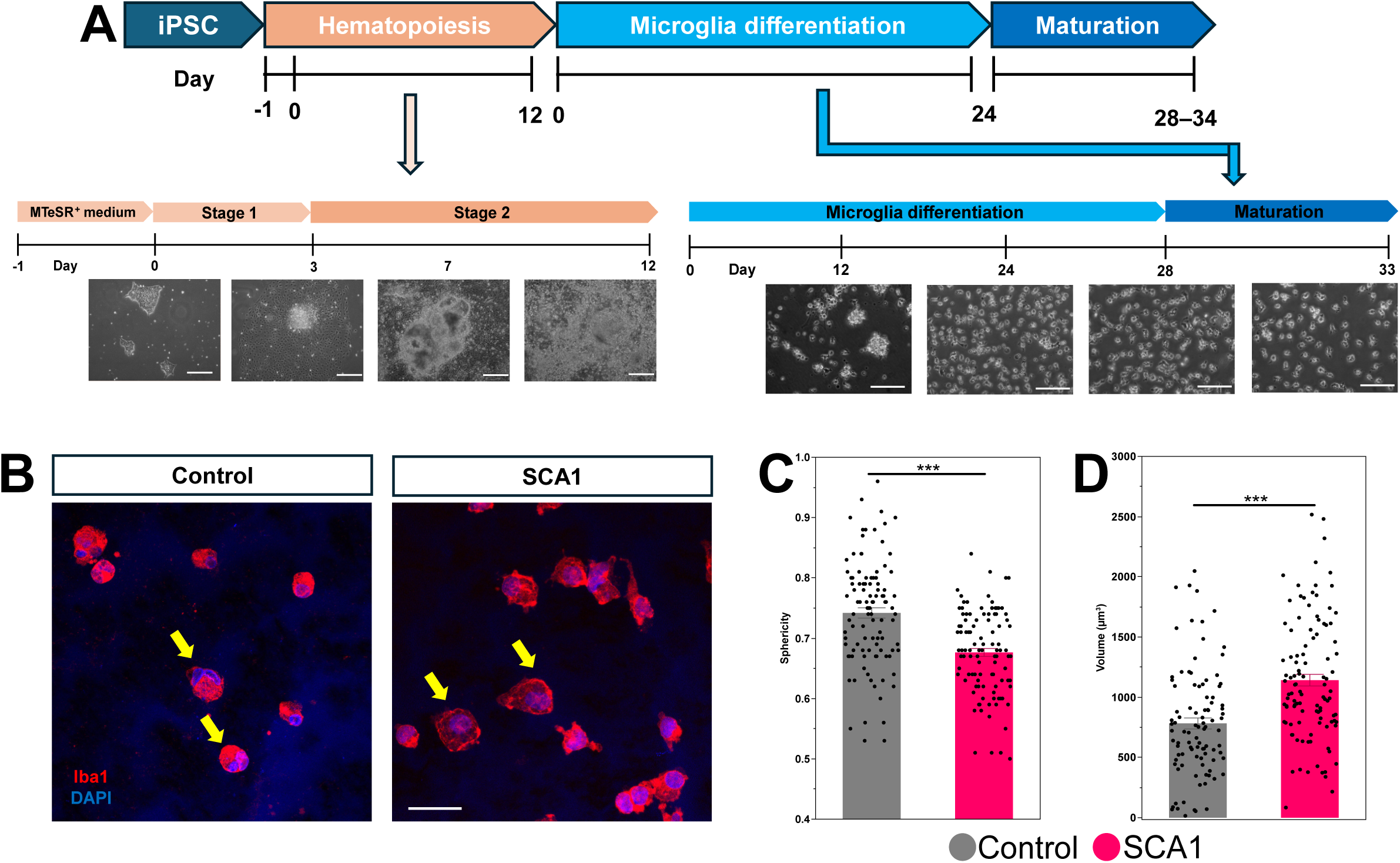
Human iPSC-derived SCA1 microglia model. A. Schematic representation of the stepwise differentiation workflow from human induced pluripotent stem cells (iPSCs) to hematopoietic progenitor cells (HPCs) and to mature iPSC-derived microglia with representative phase-contrast images showing morphological transitions from iPSCs to HPCs over the course of differentiation at day 0, 3, 7, and 12. Scale bar 100 µm. Representative phase-contrast images showing morphological transitions from HPCs to microglia over the course of differentiation at day 12, 18, and 24 as well as maturation at day 28 and 33. Scale bar 50 µm. B. Representative immunofluorescence images with Iba1 antibody showing the morphology of mature iPSC-derived SCA1 and control microglia. C-D. Imaris quantification of sphericity (C) and soma size (D).

One of the main roles of microglia is phagocytosis. The phagocytic activity of iPSC-derived microglia can be significantly increased with appropriate activation. Previous study reported that iPSC-derived microglia exhibit increased uptake of pHrodo-labeled bioparticles upon activation of hiPSC-astrocytes mediated neuroinflammatory cytokines and chemokines (Tcw et al. 2017). To investigate how *mATXN1* impacts the phagocytosis we evaluated uptake of fluorescently labeled pHrodo beads using flow cytometry (Garcia-Reitboeck et al. 2018) and IF. We found both increased pHrodo signal per SCA1 microglia and increased in the percentage of phagocytic microglia in SCA1 compared to unaffected controls (82 vs 72%) (Figure 2A-C), indicating increased phagocytosis in SCA1 human microglia.

**Figure 2.**
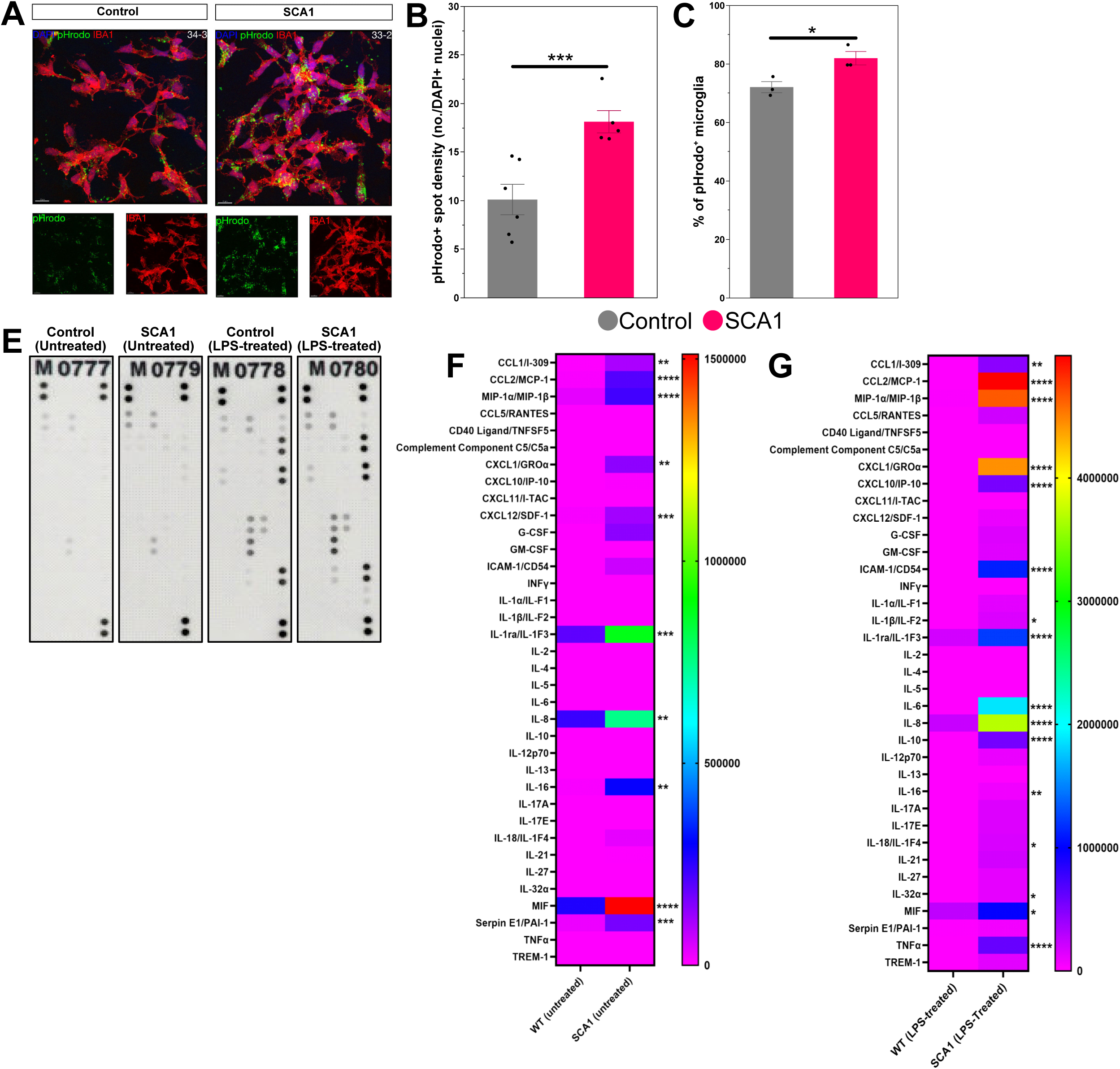
Morphological and functional characterization of human iPSC-derived microglia. A. Assessment of phagocytic capacity using pHrodo™ Green–labeled bioparticles. Fluorescent signal indicates active internalization and acidification of phagosomes in microglia. B-C. Quantification of fluorescence intensity confirms robust phagocytic activity compared to control conditions. (E-G) Cytokine array profiling of culture supernatants reveals secretion of microglia-associated cytokines and chemokines in untreated (F) and treated (G) with LPS. Data represents *n* = 3 independent iPSC lines (A–D) and *n* = 2 independent iPSC lines with 2 replicates each (E-G). Scale bars, 100 µm. Data are presented as mean ± SEM. One-way ANOVA with Tukey’s post hoc and two-tailed Welch’s t-test, * p < 0.05, *** p < 0.001.

Microglia activation is marked by the release of inflammatory cytokines. Previous studies demonstrated elevated levels of pro-inflammatory cytokines such as tumor necrosis factor (TNF), MCP-1 and interleukin-6 (IL-6) in cerebella of SCA1 mice (Aikawa et al. 2015). Cytokine microarray demonstrated an increase in inflammatory cytokines including CCL1, MIF, MCP-1 and IL16 secreted by untreated SCA1 compared to control human microglia (Figure 2E-F). We next examined whether this increased inflammatory primed state of SCA1 microglia at basal level will result in a stronger response to inflammatory stimuli. Inflammatory stimulation with LPS (100 ng/ml) induced secretion of the proinflammatory cytokines including TNFα, MCP-1 and IL-6 that were significantly larger in SCA1 microglia (Figure 2G).

Together these results indicate that *mATXN1* expression is sufficient to cause pro-inflammatory phenotype in human microglia, increasing their soma size, phagocytosis and pro-inflammatory cytokine secretion in a cell-autonomous manner.

### Mutant ATXN1 alters expression of genes and proteins in human microglia

Previous studies indicated that ATXN1 plays a role in regulating transcription and that nuclear localization of mATXN1 is key for SCA1 pathogenesis (Serra et al. 2004; Handler et al. 2023). To determine how mATXN1 impacts gene expression in human microglia, we performed RNA sequencing of SCA1 and unaffected control human microglia. We found a total of 426 DEG (p<0.05, Log2FC > 0.6 or < -0.6), with 228 downregulated and 198 upregulated DEGs (Figure 3A). Among downregulated genes are *CCL2, TCEF1* and *MYO10* that play a role in inflammation, protection from STING activation and phagocytosis. Among the upregulated genes are many mitochondrial genes including MT-AT8, *MT-CO2,* and *MT-ND4L*, involved in oxidative phosphorylation, *SCARNA7* and *DDX52*, involved in RNA processing, *CHRNA1*, acetylcholine receptor, potassium channel KCNJ5 and dual specificity phosphatase *DUSP16* shown to promote M2 microglial phenotype.

**Figure 3.**
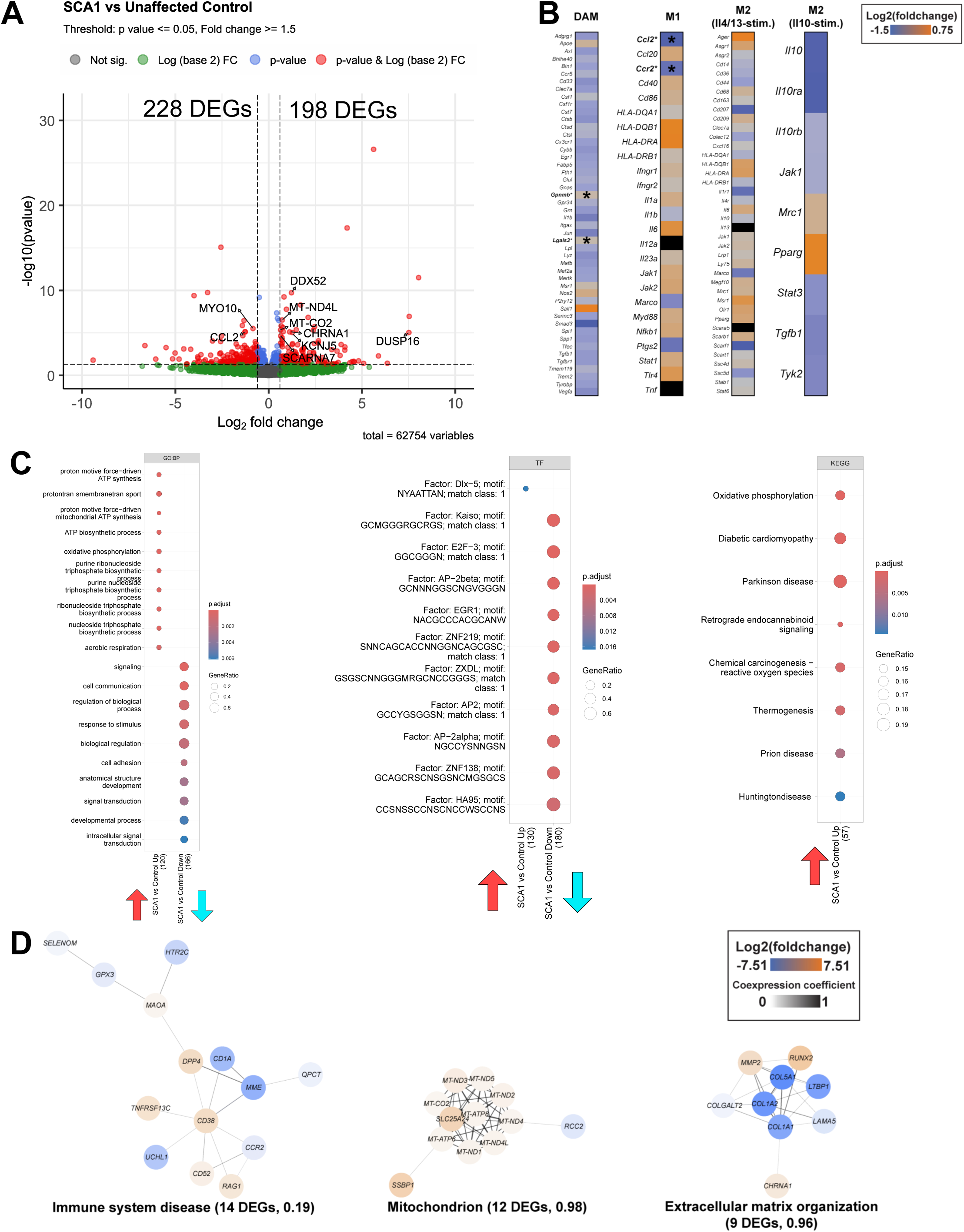
**Mutant ATXN1 alters transcription in human SCA1 microglia**. RNA was isolated from day 30 microglia derived from SCA1 and unaffected iPSCs and RNA seq were performed. A. RNA seq Volcano plots. B. Altered expression of DAM, M1 and M2 genes in SCA1 microglia. C. Gene Ontology pathways of SCA1 microglia upregulated genes in biological processes, cellular components, and molecular function. D. Upregulated SCA1 microglia transcription factors.

To provide insight into the microglia phenotype, we examined expression of M1, M2 and DAM genes in SCA1 human microglia DEGs. We have found a spectrum of expression of these genes, with an overall decrease in DAM genes and both increased and decreased expression of M1 and M2 genes indicating that mATXN1 expression induces a reactive phenotype characterized by a mixed expression of M1 and M2 microglia genes (Figure 3B). Compared with previously published data, we found 34 DEGs that are altered both in our dataset and in human SCA1 microglia in single-nuclei RNA seq of postmortem human SCA1 cerebellum, and of 20 DEGs altered in our data and in human SCA1 motor neurons (MNs) (Supplementary Figure 3A and C). Interestingly, we found a larger overlap of transcriptional factors (TF) with over 2/3 and 1/3 of TF involved in SCA1 microglia also playing role in microglia from SCA1 patient cerebella and SCA1 MNs (Supplementary Figure 3B and D). This result may indicate that mATXN1 alters transcription differently in different cell types or disease stages, but that basic molecular mechanism remains similar.

To gain insight into how these DEGs impact microglia we performed pathway analysis (Figure 3C). GO analysis of upregulated genes identified pathways associated with mitochondrial function including proton transmembrane transporter activity (GO:MF p=5.3 X 10-7), proton motive force driven mitochondrial ATP synthesis (GO:BP p= 5.3 X 10-6) and respiratory chain complex (GO:CC p= 1.6X10-6). KEGG pathway analysis identified oxidative phosphorylation (p=1.5 X10-5), Huntington disease (p= 1.2 X10-2) and ALS (p=0.4 X 10-2). Transcription factor analysis identified DLX5 and HOXA2 as the main transcriptional regulators of upregulated DEGs.

Pathways impacted by downregulated genes were protein binding (GO:MF p=3.8X 10-5), signaling (GO: BP p=3.2 X10-5), endomembrane system (GO CC p= 3.3 10-5) and extracellular matrix (GO:CC p= 3.4 X 10-2). Among the transcription factors identified were Kaiso (p =1 X 10-5), E2F3 (p =4.2 X 10-5) and AP-2 (p= 1.9 X 10-4). Overall, the three main pathways that were altered in SCA1 microglia were the immune system, mitochondrion and extracellular matrix (Figure 3D).

Proteomic analysis of human SCA1 and control microglia identified 185 differentially expressed proteins (DEPs) with 72 upregulated and 113 downregulated proteins (Figure 4A). Among the upregulated proteins are mitochondrial proteins including MT-CO2, also detected in our transcriptome, ATP5MJ, COXC7, NDUFA9, NDUFB8, and NDUFA12.

**Figure 4.**
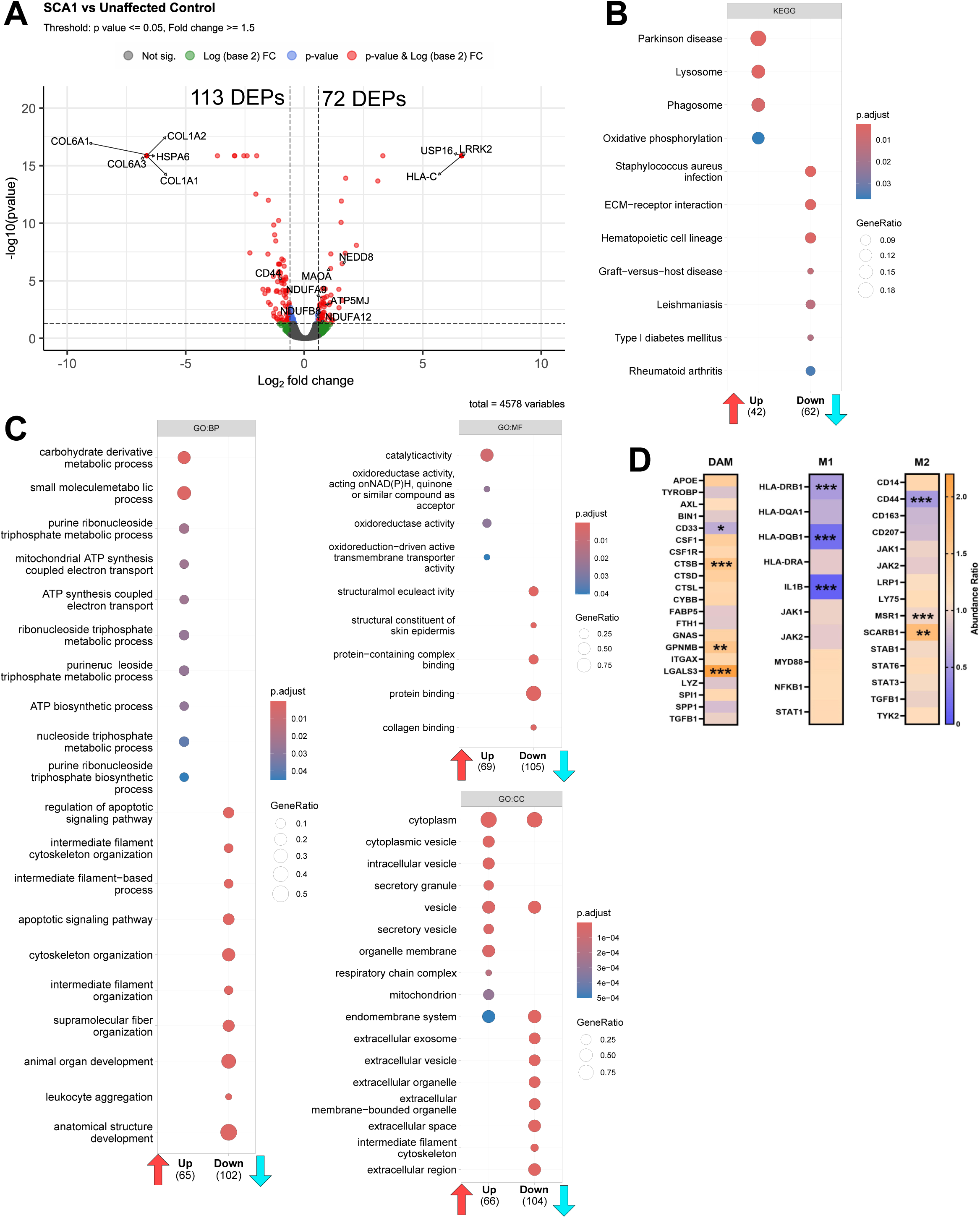
Mutant ATXN1 expression alters protein expression in human SCA1 microglia. A. Volcano plot of human SCA1 vs unaffected control proteomics. B. Kyoto encyclopedia of genes and genomes pathways of SCA1 microglia proteins. C. Gene ontology pathways of SCA1 microglia proteins in biological processes, molecular functions, and cellular components. D. Altered levels of DAM, M1 and M2 proteins in SCA1 microglia.

Intriguingly, we found significant increase in the expression of Ubiquitin carboxyl-terminal hydrolase 16 (USP16), leucine-rich repeat kinase 2 (LRRK2), monoamine oxidase A (MAOA), and NEDD8. As these proteins contribute to microglia activation, increase in their expression may underlie increased reactive phenotype of SCA1 microglia. In addition, USP16, MAOA and NEDD8 all contribute to NF-kB activation, providing the possible mechanism for the increased reactivity of SCA1 microglia. Expression of antigen presenting proteins HLA-C and HLA-DM is also increased in SCA1 microglia further supporting reactive phenotype. Increased expression of proteins involved in phagocytosis, including CDC42 involved in phagocytic cup formation, and cathepsin B, involved in phagocytosis of apoptotic neurons, may underlie increased phagocytosis by SCA1 microglia.

Consistent with the transcriptomic data, expression of many extracellular matrix (ECM) proteins was downregulated including COL1A1, COL1A2, COL6A1, COL6A3, CD44. In addition, expression of heat shock protein HSPA6 (Hsp70), previously shown to ameliorate pathogenesis in SCA1 mouse model, was significantly decreased in SCA1 microglia. Intriguingly, protein levels of both STING1 and IL1B, increased in many neurodegenerative diseases, are significantly decreased in SCA1 microglia.

Among pathways identified using downregulated proteins were structural molecule activity (GO:MF p= 5.3 10-11), cytoskeletal protein, collagen and actin binding (GO:MF p= 2,7X 10-4, 3.3 X10-2 and 3.4 X10-2 respectively), intermediate filament cytoskeletal organization (GO:BP p = 5.3 X 10-7), extracellular exosome, vesicle and space (GO:CC p = 1.3 X10-17, p = 2.4 X10-17, p=3.6 X10-15) (Figure 4 B-C). Pathways associated with increased protein expression included catalytic activity ( GO:MF p= 1.5 10-4), carbohydrate derivative metabolic process (GO:BP p=1.3 X10-3), phagocytosis (GO:BP p= 2.8 X10-2), mitochondrial ATP synthesis (GO:BP p =3.5X 10-2), cytoplasm (GO:CC p =1.5 X10-11), secretory vesicle (GO:CC p =3.7X10-7), phagosome (KEGG p= 1.5.9 X X10-3), lysosome (KEGG p =5.9X10-3) (Figure 4B-C). These results provide mechanism for the increased in phagocytosis and inflammatory cytokine secretion in SCA1 microglia and provide support for the cell-autonomous effects of mutant ATXN1 in microglia.

Together our results indicate that mutant ATXN1 expression induces significant changes in expression of genes and proteins indicating alterations in metabolism, reactivity, phagocytosis and extracellular matrix.

### Cell-autonomous effects of mutant ATXN1 on microglial phenotype in SCA1 mice

To determine how expression of mutant ATXN1 impacts microglial phenotype in SCA1 when other cells also contribute to microglia activation, we used a novel conditional SCA1 model, *f-ATXN1^146Q/2Q^* mice. In these mice, coding regions of human *ATXN1* with 146 CAG repeats are surrounded by LoxN sites and knocked into one of the endogenous mouse *Atxn1* allele (Duvick et al. 2024) allowing cell-type specific deletion of *mATXN1*. We generated SCA1 microglial knock out mice (*ATXN1_mKO_*), by crossing *f-ATXN1^146Q/2Q^* and *Lyve1^CRE^* mice (Figure 6A). In these mice, Cre recombinase is expressed under control of the lymphatic vessel endothelial hyaluronan receptor 1+ (*Lyve1*) promoter in erythro-myeloid progenitor cells, which become microglia, and macrophages (Pham et al 2010). We used *Lyve1^CRE^* mice because previous studies found very effective Cre recombination (80%) in microglia (Scott et al 2022). We isolated microglia using magnetic beads and using RT-qPCR of enriched microglia (Supplementary Figure 4) we found a 60% reduction of *mATXN1* in *ATXN1_mKO_* microglia as compared to *f-ATXN1^146Q/2Q^* microglia.

To determine how microglial *mATXN1* expression impacts microglia density and morphology in SCA1 cerebella we used IF with Iba1, confocal microscopy and image analysis. As previously shown in other mouse models we found increased microglial density in the cerebellar cortex of 35-week-old *f-ATXN1^146Q/2Q^* mice that was ameliorated in *ATXN1_mKO_* mice (Figure 5B). We used Imaris 3D reconstructions to further characterize microglial morphological changes in SCA1 mice. Intriguingly, we have not found any significant changes in microglial processes, soma size and branching in *f-ATXN1^146Q/2Q^* microglia compared to WT mice. More surprisingly, we found a reduction in microglial processes, soma size and branching in *ATXN1_mKO_* mice compared to both *f-ATXN1^146Q/2Q^* and WT mice (Figure 5D-I), indicating that expression of mutant ATXN1 in microglia is required to maintain their morphology in SCA1.

**Figure 5.**
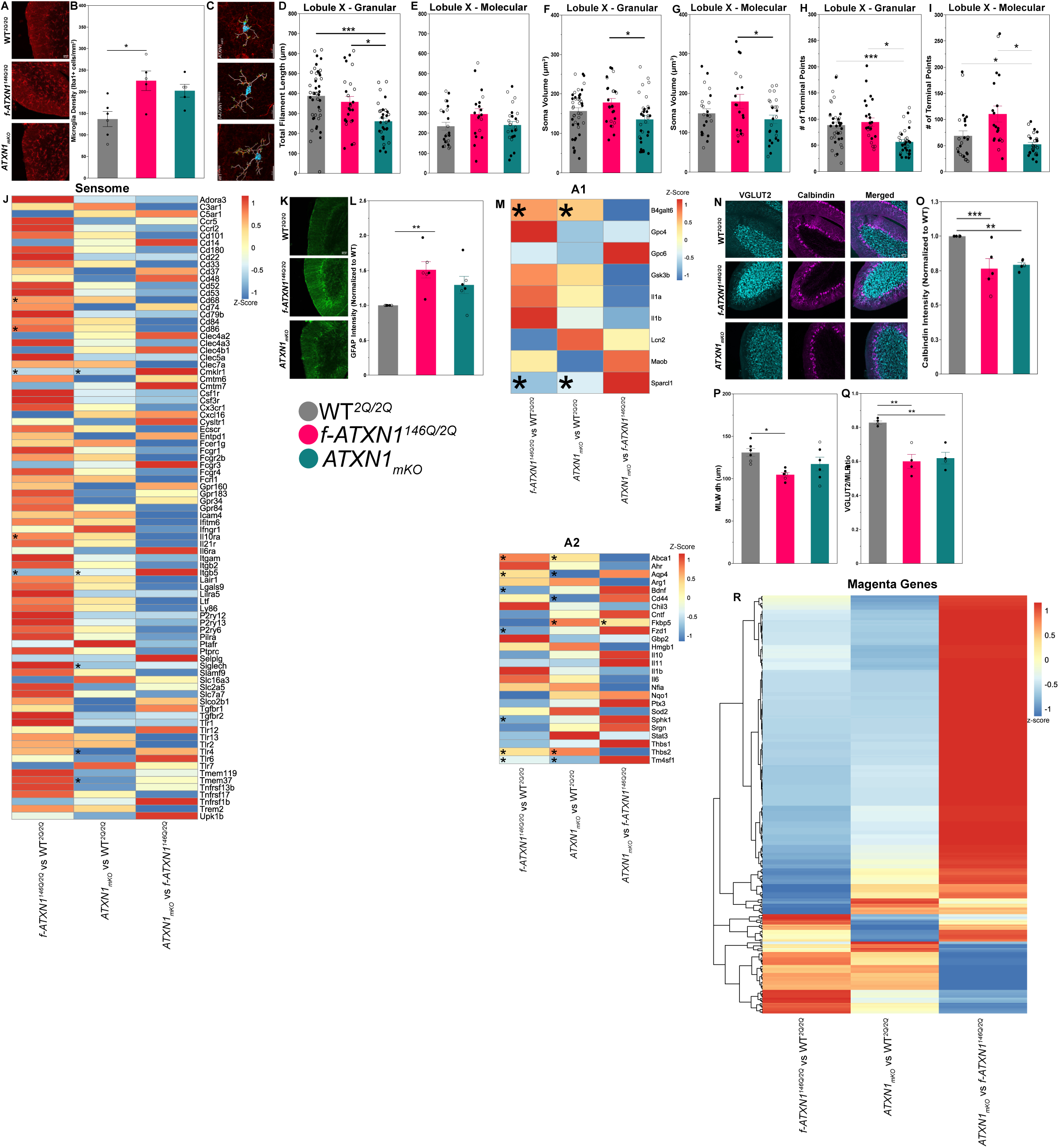
Impact of selective deletion of mutant ATXN1 on microglia reactivity and cerebellar pathogenesis. A. Representative images of Iba1+ cells in 34-week mouse lobule X. B. Quantification of microglia density. N=5 WT*^2Q/2Q^*, N=5 *f-ATXN1^146Q/2Q^*, and N=5 *ATXN1_mKO_* mice 34–36 weeks of age. C. Representative Imaris 3D reconstruction images in cerebellar lobule X. D. Lobule X granular layer total filament length. E. Lobule X molecular layer total filament length. F. Lobule X granular layer soma volumes. G. Lobule X molecular layer soma volumes. H. Lobule X granular layer number of terminal points. I. Lobule X molecular layer number of terminal points. J. Heatmap of sensome genes. Measured using z-score. K. Representative images of GFAP intensity in 32–35-week-old mouse lobule X. L. Quantification of GFAP intensity. N=6 WT*^2Q/2Q^*, N=6 *f-ATXN1^146Q/2Q^*, and N=6 *ATXN1_mKO_* mice 32-–35 weeks of age. M. Heatmaps of 35-week cerebellar A1 gene expression (*Top*) and A2 gene expression (*Bottom*). N. Representative images of VGLUT2, Calbindin, and merged intensities. O. Quantification of Calbindin intensity. N=5 WT*^2Q/2Q^*, N=5 *f-ATXN1^146Q/2Q^*, and N=5 *ATXN1_mKO_* mice 32-35 weeks of age. P. Quantification of molecular layer width. N=6 WT*^2Q/2Q^*, N=6 *f-ATXN1^146Q/2Q^*, and N=6 *ATXN1_mKO_* mice 32–35 weeks of age. Q. Quantification of VGLUT2 width over molecular layer width. N=4 WT*^2Q/2Q^*, N=4 *f-ATXN1^146Q/2Q^*, and N=4 *ATXN1_mKO_* mice 32–35 weeks of age. R. 35-week cerebellar magenta gene expression. 214 genes that were significantly altered in *f-ATXN1^146Q/2Q^* and *ATXN1_mKO_* as compared to WT. Data is average ± SEM with individual mice presented as dots. Open dots represent female mice while filled dots represent male mice. One-way ANOVA with Tukey’s test. * P < 0.05, ** P < 0.01, *** P < 0.001.

**Figure 6.**
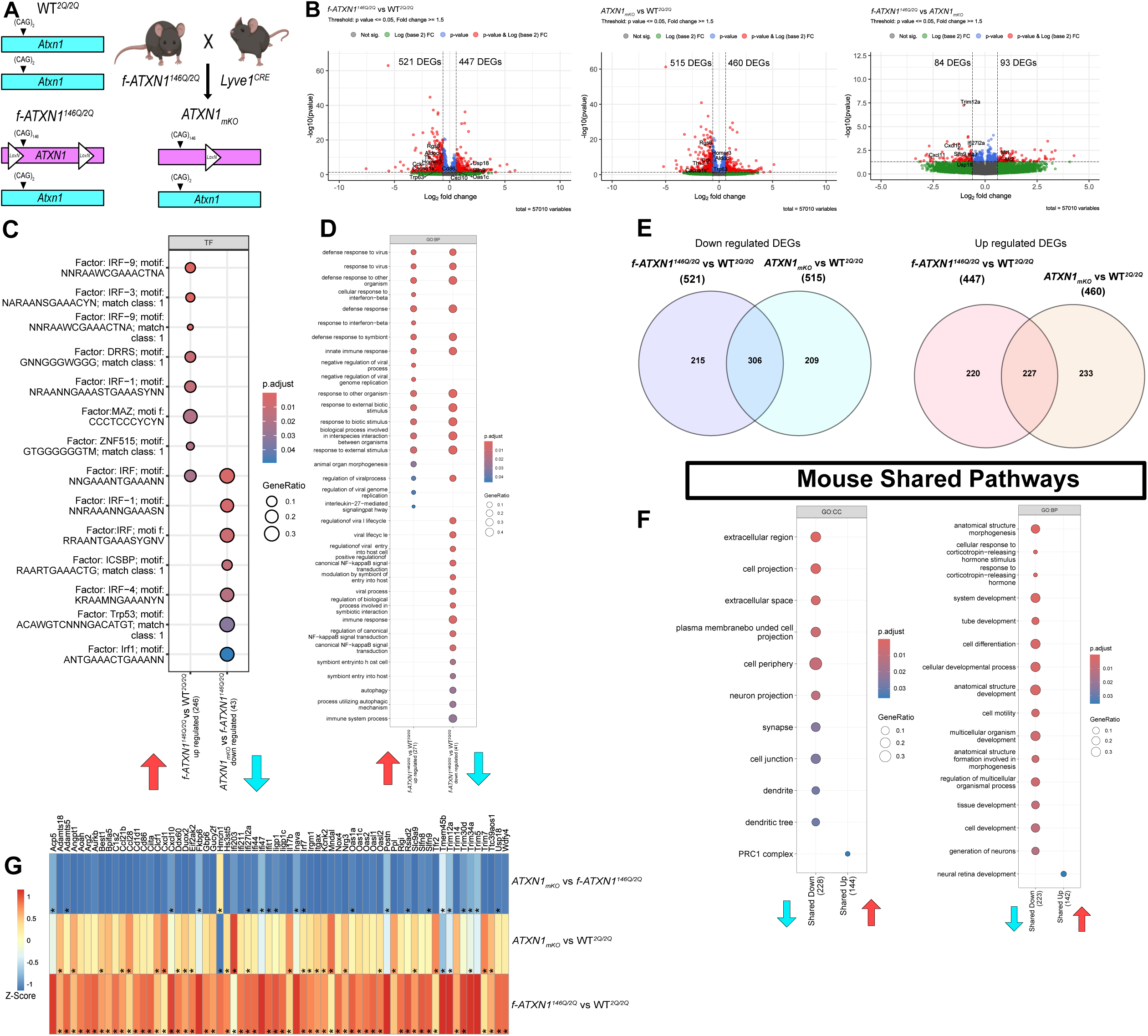
Selective deletion of mutant *ATXN1* in microglia of conditional SCA1 mice ameliorates IFN pathway. A. Schematic of mouse creation. B. RNA sequencing volcano plots. *Left*, *f-ATXN1^146Q/2Q^* vs WT*^2Q/2Q^* comparison. *Middle*, *ATXN1_mKO_* vs *f-ATXN1^146Q/2Q^* comparison. *Right*, *ATXN1_mKO_* vs *f-ATXN1^146Q/2Q^* comparison. C. Gene Ontology Biological Processes pathway analysis. Left column contains upregulated pathways in *f-ATXN1^146Q/2Q^* vs WT*^2Q/2Q^* comparison, right column contains downregulated pathways in *ATXN1_mKO_* vs *f-ATXN1^146Q/2Q^* comparison. D. Transcription factor analysis. Left column contains upregulated pathways in *f-ATXN1^146Q/2Q^* vs WT*^2Q/2Q^* comparison, right column contains downregulated pathways in *ATXN1_mKO_* vs *f-ATXN1^146Q/2Q^* comparison. E. Venn diagram of DEGs. *Left*, down regulated DEGs. 521 DEGs in *f-ATXN1^146Q/2Q^* vs WT*^2Q/2Q^* comparison (blue), 515 *ATXN1_mKO_* vs WT*^2Q/2Q^* comparison (cyan). *Right*, up regulated DEGs. 447 DEGs in *f-ATXN1^146Q/2Q^*vs WT*^2Q/2Q^* comparison (red), 460 *ATXN1_mKO_* vs WT*^2Q/2Q^* comparison (orange). F. Shared pathways. *Left*, Gene Ontology Cellular Component pathway analysis. *Right*, Gene Ontology Biological Processes pathway analysis. G. Heatmap of immune response genes. Measured using z-score.

As our results indicate a reduced ramification in *ATXN1_mKO_* mice and one important role of microglial ramification is sensing the local environment through the sensome (Hickman et al. 2013; 2018) we examined the expression of sensome genes. Consistent with reduced ramification, we found a decreased expression of sensome genes in *ATXN1_mKO_* mice compared to *f-ATXN1^146Q/2Q^* mice (Figure 5H). Taken together this data indicates a significant contribution of microglial *mATXN1* to microglial morphology and sensing in SCA1.

### Microglial mutant *ATXN1* expression contributes to cerebellar pathogenesis

Next, we assessed the contribution of microglial ATXN1 to cerebellar pathogenesis using IF with calbindin, a marker for Purkinje cell health decreased in SCA1, and glial fibrillary acidic protein (GFAP) marker of astrogliosis increased in SCA1.

While GFAP expression was increased in *f-ATXN1^146Q/2Q^* mice it was not significantly altered in *ATXN1_mKO_* mice (Figure 5L). Furthermore, to determine the neurotoxicity state of astrocytes we assessed A1 and A2 gene expression. We found that *f-ATXN1^146Q/2Q^* astrocytes exhibit increased A1 gene expression and mixed A2 gene expression when compared to WT, and *ATXN1_mKO_* astrocytes show ameliorated expression of A1 and A2 genes when compared to *f-ATXN1^146Q/2Q^* (Figure 5M), indicating the role of microglial ATXN1 in cerebellar astrogliosis and astrocyte-microglia communication Similarly, PC molecular layer width, measure of dendritic atrophy, was statistically reduced only in *f-ATXN1^146Q/2Q^* (Figure 5P). Moreover, *f-ATXN1^146Q/2Q^* mice exhibit a dominantly downregulated expression of magenta gene, an indicator of worsened PC health in SCA1 (Ingram et al., 2016) when compared to WT, while *ATXN1_mKO_* exhibit ameliorated expression of magenta genes when compared to *f-ATXN1^146Q/2Q^* (Figure 5R). Reduced PC calbindin intensity, another measure of PC health, and a reduction in climbing fiber synapses on PC (measured as VGLUT2/ML ratio) are indistinguishable in *f-ATXN1^146Q/2Q^* and *ATXN1_mKO_* mice (Figure 5O, Q), indicating that mATXN1 in microglia does not contribute to these aspects of PC pathology in SCA1 mice.

These findings indicate that microglial mATXN1 expression contributes to cerebellar astrogliosis and aspects of PC pathology in SCA1 mice.

### Microglial mutant *ATXN1* increases mouse viral response gene expression

To determine how microglial *mATXN1* impacts altered gene expression in SCA1 cerebellum we performed bulk RNA sequencing of cerebella dissected from 35-38-week-old WT, *f-ATXN1^146Q/2Q^* and *ATXN1_mKO_* mice. We compared gene expressions in *f-ATXN1^146Q/2Q^* vs WT, *ATXN1_mKO_* vs WT, and *ATXN1_mKO_* vs *f-ATXN1^146Q/2Q^*(Figure 6).

We found 968 DEGs (P < 0.05, Log2FC < -0.6 or > 0.6) in *ATXN1^146Q/2Q^* with 521 downregulated and 447 upregulated genes. Genes downregulated in *f-ATXN1^146Q/2Q^* mice include genes previously identified in SCA1 including *Homer3*, *Rgs8*, *Aldoc* previously identified as related to SCA1 Purkinje cell health (Ingram et al. 2016), *Cacna1s* and *Th* involved in synapses, and *Cck* a gene known to be protective in SCA1 (Wozniak et al. 2021). Among upregulated DEGs were *Cxcl10*, *Oas1c*, *Slfn9*, *Usp18*, *Cd86* known to be involved in immune response and inflammation (Majumdar et al. 2025; P. Zhang et al. 2024) (Figure 6B, *left panel*). Pathway analysis identified increase in viral response pathways in SCA1 cerebella including: defense response to virus (GO:BP P = 1.1 X10-6), response to virus (GO:BP P = 1.5 X10-6), defense response to other organism (GO:BP P = 2.2 X10-5), defense response (GO:BP P = 2.5 X10-4), defense response to symbiont (GO:BP P = 3.5 X10-4), and innate immune response (GO:BP P = 7.7 X10-4), response to other organism (GO:BP P = 1.9 X10-3), response to external biotic stimulus (GO:BP P = 2.0 X10-3), response to biotic stimulus (GO:BP P = 3.7 X10-3), biological process involved in interspecies interaction between organisms (GO:BP P = 4.6 X10-3), response to external stimulus (GO:BP P= 6.7 X10-3), and regulation of viral process (GO:BP P = 4.1 X10-2) (Figure 6D). Consistent with this, transcription factor analysis identified TF involved in interferon response: IRF-9 (motif: NNRAAWCGAAACTNA, P = 2.2 X10-6), IRF-3 (motif: NARAANSGAAACYN, P= 1.7 X10-5), and IRF-1 (motif: NRAANNGAAASTGAAASYNN, P= 1.2X10-2) (Figure 6C). *ATXN1_mKO_* vs WT*^2Q/2Q^* comparison revealed 975 DEGs with 515 downregulated and 460 upregulated genes (Figure 6B *middle panel*). Importantly, we found 177 DEGs comparing *ATXN1_mKO_* vs *f-ATXN1^146Q/2^*^Q^ cerebella with 84 downregulated and 93 upregulated DEGs. Among genes downregulated in *ATXN1_mKO_* compared to *f-ATXN1^146Q/2Q^* mice were *Trim12a*, *Trim5*, and *Trim30d* involved in defense response and positive regulation of NF-kB transcription factor activity (L. Zhang et al. 2024; Tareen et al. 2009; Bieniasz 2004), *Trim34a* and *Tmem45b* involved in innate immune response (Vidal-Itriago et al. 2022; Yan et al. 2022), and *Ifi47*, *Ifi28l2a*, *Iigp1*, and *Ifit1* interferon inducible genes involved in defense response (Gilly et al., n.d.; Kim et al. 2025; Liesenfeld et al. 2011; Kimura et al. 2013). Pathway analysis of downregulated DEGs identified defense response to virus (GO:BP P = 3.7 X10-5), response to virus (GO:BP P = 1.20944E-5), defense response to other organism (GO:BP P =5.3 X10-7), defense response (GO:BP P =8.1 X10-5), defense response to symbiont (GO:BP P =7.0 X10-6), innate immune response (GO:BP P =2.9 X10-6), response to other organism (GO:BP P =6.2 X10-7), response to external biotic stimulus (GO:BP P =6.5 X10-7), response to biotic stimulus (GO:BP P =9.2 X10-7), biological process involved in interspecies interaction between organisms (GO:BP P =1.9 X10-6), response to external stimulus (GO:BP P=3.5 X10-5), and regulation of viral process (GO:BP P =4.4 X10-6). Transcription factor analysis uncovered downregulation in the interferon response transcription factor IRF (motif: NNGAAANTGAAANN, P = 4.2 X10-3), IRF-1 (motif: NNRAAANNGAAASN, P = 5.5 X10-3), IRF-4 (motif: KRAAMNGAAANYN P = 1.8 X10-2), and IRF (motif: RRAANTGAAASYGNV P = 8.8 X10-3) (Figure 7C). These findings indicate that microglial mutant ATXN1 contributes to increased innate immune and interferon responses in SCA1 cerebella (Figure 6B *right panel* and 6G).

**Figure 7.**
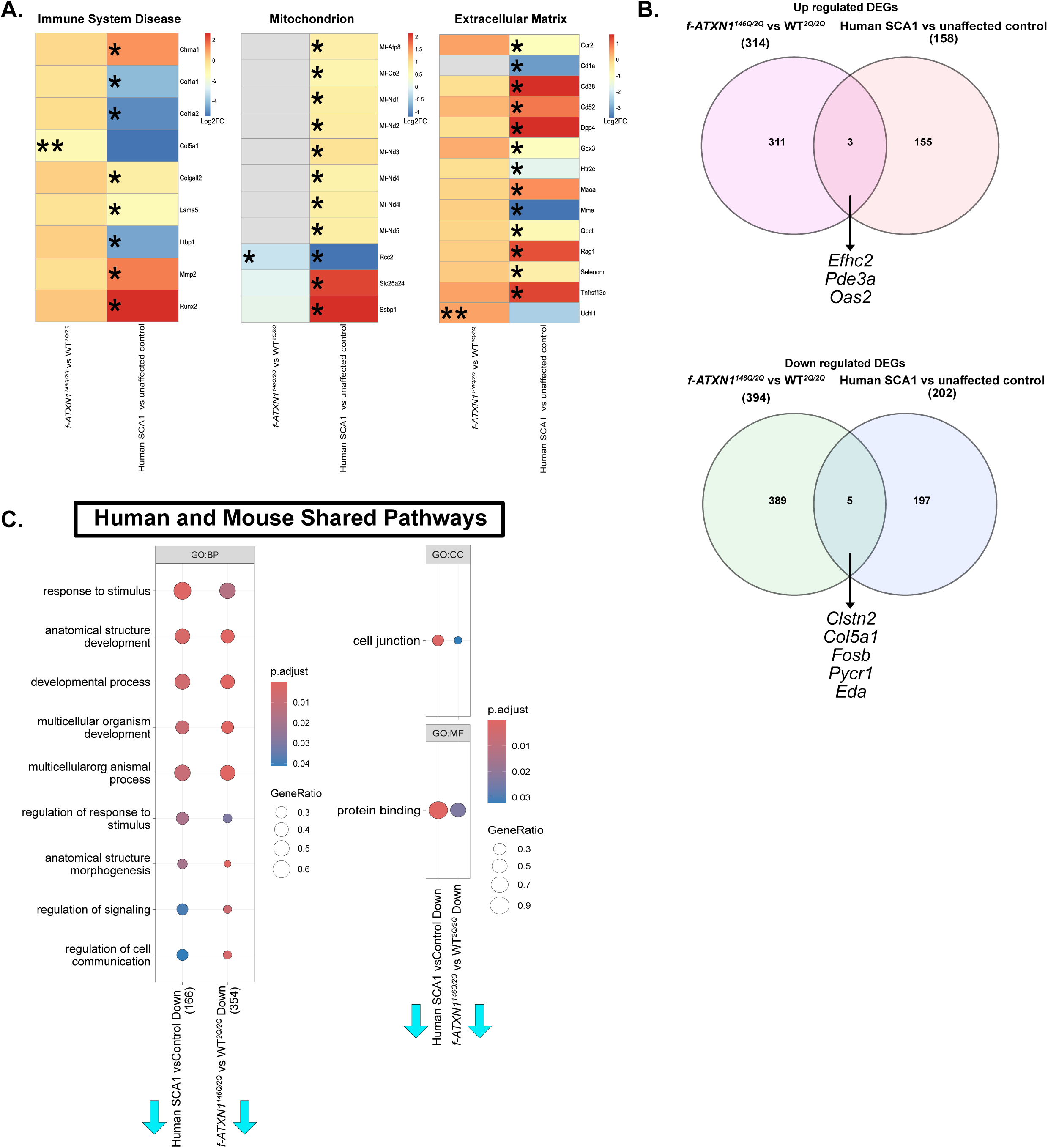
Comparing human to mouse transcriptomics reveals differences in DEGs. A. Heatmaps using log2FC. *Left*, immune system disease genes. *Middle*, mitochondrion DEGs. *Right*, extracellular matrix DEGs. B. Venn diagrams of DEGs in each comparison. *Top*, up regulated DEGs. 314 DEGs *f-ATXN1^146Q/2Q^* vs WT*^2Q/2Q^* comparison (pink). 158 DEGs Human SCA1 vs unaffected controls (orange). *Bottom*, down regulated DEGs. 394 DEGs *f-ATXN1^146Q/2Q^* vs WT*^2Q/2Q^* comparison (green). 202 DEGs Human SCA1 vs unaffected controls (blue). C. Shared pathways. *Left*, Gene Ontology Biological Processes pathway analysis. *Right,* Gene Ontology Cellular Component pathway analysis.

To determine genes and pathways that are less influenced by the microglial *mATXN1*, we compared expression of down regulated and up regulated genes between *f-ATXN1^146Q/2Q^* vs WT and *ATXN1_mKO_* and WT comparisons and found shared 306 downregulated and 227 upregulated DEGs (Figure 7F). Furthermore, when assessing the shared pathways we find a shared down regulated pathways including: extracellular region (GOCC P = 3.7 X 10-4), extracellular space (GOCC P = 3.8 X 10-3), cell junction (GOCC P = 2.6 X 10-2), synapse(GOCC P = 2.4 X 10-2), dendrite(GOCC P = 2.8 X 10-2), and dendritic tree (GOCC P = 2.9 X 10-2), system development (GOBP P = 4.7 X 10-4), cell differentiation (GOBP P = 2.1 X 10-3), cellular developmental process (GOBP P = 2.1 X 10-3), cell development (GOBP P = 9.6 X 10-3), tube development (GOBP P = 8.5 X 10-4). These results indicate that *mATXN1* in microglia significantly contributes to interferon related immune response genes but is less involved in development and synapse related transcriptional alterations.

Finally, we compared gene expression changes in human SCA1 microglia to genes altered in *ATXN1^146Q/2Q^* mice as having a humanized version of *mATXN1* allows for a more direct comparison between human and mouse tissue. Comparing DEGs between human microglia and mouse cerebella reveals only 3 shared up regulated and 5 shared down regulated genes (Figure 7B). Shared up regulated genes include: *Efhc2*, involved in epilepsy and Turner syndrome, *Pde3a*, involved in cardiovascular function, and *Oas2*, known to activate type I interferon pathways to inhibit viral replication (Liao et al. 2020). Shared down regulated genes include: *Clstn2*, involved in synapse assembly, *Col5a1*, involved in the extracellular matrix, *Fosb*, transcription activator, *Pyrc1*, involved in mitochondrial function, and *Eda*, involved in cell-cell signaling. Furthermore, we find only shared down regulated pathways (Figure 7C). These pathways include: response to stimulus (GOBP, human P = 7.9 X 10-4, mouse P = 1.5 X 10-2), anatomical structure development (GOBP, human P = 2.9 X 10-3, mouse P = 8.6 X 10-7), developmental process (GOBP, human P = 5.8 X 10-3, mouse P = 4.25 X 10-6), multicellular organism development (GOBP, human P = 8.5 X 10-3, mouse P = 6.6 X 10-7), multicellular organismal process (GOBP, human P = 9.1 X 10-3, mouse P = 7.8 X 10-5), regulation of response to stimulus (GOBP, human P = 1.8 X 10-2, mouse P = 3.0 X 10-2), anatomical structure morphogenesis (GOBP, human P = 2.0 X 10-2, mouse P = 2.2 X 10-6), regulation of signaling (GOBP, human P = 3.9 X 10-2, mouse P = 7.3 X 10-3), regulation of cell communication (GOBP, human P = 4.1 X 10-2, mouse P = 3.9 X 10-3), cell junction (GOCC, human P = 1.5 X 10-3, mouse P = 3.2 X 10-2), and protein binding (GOMF, human P = 3.8 X 10-5, mouse P = 2.3 X 10-2). Importantly, three main pathways that were altered in SCA1 human microglia (Figure 3C), mitochondria, the immune system disease and extracellular matrix were also altered in SCA1 mice (Figure 7A). Taken together these data point towards similarities and differences between SCA1 microglia in human and mouse models of disease.

### Microglial mutant *ATXN1* expression contributes to motor, cognitive, and mood deficits in SCA1 mice

To determine how microglial mATXN1 expression impacts SCA1 behavioral phenotypes, we performed a battery of motor, cognitive and mood assays.

Rotarod is a gold standard for assessing motor coordination in SCA1 mouse models (Clark et al. 1997; Watase et al. 2002; Duvick et al. 2024). Previous work shows *f-ATXN1^146Q/2Q^* also exhibit a reduced latency to fall time (Duvick et al. 2024). To assess progressive decrease in motor coordination we used rotarod at 14, 24, and 34 weeks. Both *f-ATXN1^146Q/2Q^*and *ATXN1_mKO_* mice display reduced latency to fall times compared to WT at 14 and 24 weeks. However, at 34 weeks, *ATXN1_mKO_* mice show significantly increased latency to fall compared to *f-ATXN1^146Q/2Q^* (Figure 8A), indicating ameliorated coordination deficits at the late disease stage. We also assessed balance using a balance beam, an assay which assesses foot slips while mice attempt to cross beams. At 24-weeks only *f-ATXN1^146Q/2Q^* mice had an increased number of slips on the hard square beam compared to WT mice, while performance of *ATXN1_mKO_*mice was similar to WT mice (Figure 8B). These results indicate that removing microglial mATXN1 expressions ameliorates motor deficits at late stages of disease in SCA1 mice.

**Figure 8.**
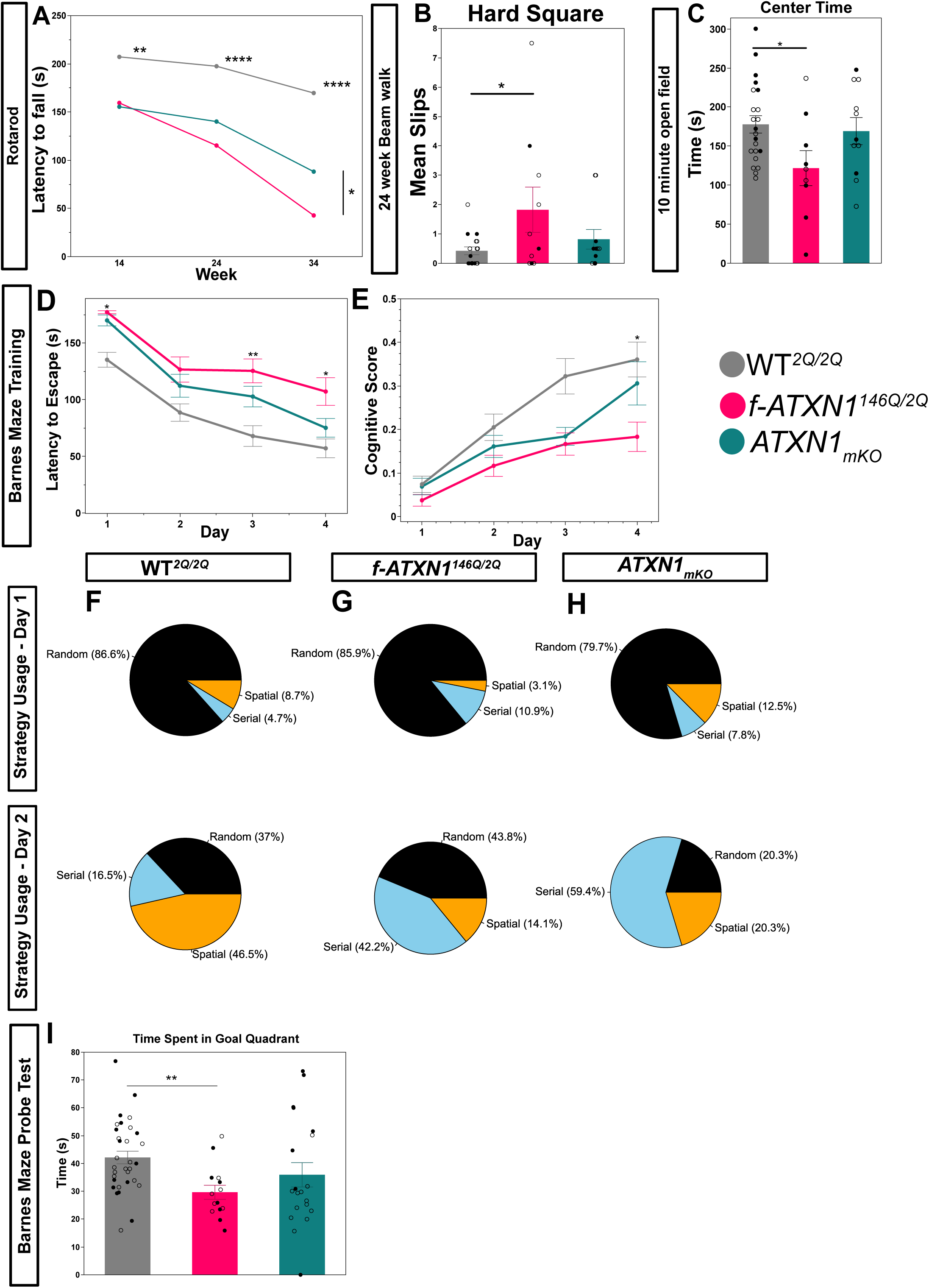
Improved performance on motor, cognitive and anxiety tests in *ATXN1_mKO_* mice. A. Rotarod latency to fall on day 4 averages across weeks. Week 14, N=30 WT*^2Q/2Q^*, N=13 *f-ATXN1^146Q/2Q^*, N=15 *ATXN1_mKO_*. Week 24, N=29 WT*^2Q/2Q^*, N=13 *f-ATXN1^146Q/2Q^*, N=15 *ATXN1_mKO_*. Week 34, N=29 WT*^2Q/2Q^*, N=13 *f-ATXN1^146Q/2Q^*, N=13 *ATXN1_mKO_*. Two-way ANOVA with Tukey’s post hoc test. B. Average number of slips during 24-week beam walk, N=18 WT*^2Q/2Q^*, N=10 *f-ATXN1^146Q/2Q^*, N=10 *ATXN1_mKO_* mice 24 weeks of age. One-way ANOVA with Tukey’s post hoc. C. Total time spent in the center zone during the first 10 minutes in the open field. N=22 WT*^2Q/2Q^*, N=9 *f-ATXN1^146Q/2Q^*, N=11 *ATXN1_mKO_* mice 13 weeks of age. One-way ANOVA with Tukey’s post hoc. D. Barnes maze latency to find the escape over the four training days. Two-way ANOVA with Tukey’s post hoc. E. Cognitive score data derived by providing BUNs software mouse movement traces during the Barnes maze training days for each mouse. Data is averaged per genotype. Two-way ANOVA with Tukey’s post hoc. F. WT*^2Q/2Q^* strategy analysis day 1 (*top*) and day 4 (*bottom*). G. *f-ATXN1^146Q/2Q^* strategy analysis day 1 (*top*) and day 4 (*bottom*). H. *ATXN1_mKO_* strategy analysis day 1 (*top*) and day 4 (*bottom*). N=32 WT*^2Q/2Q^*, N=15 *f-ATXN1^146Q/2Q^*, N=19 *ATXN1_mKO_* mice 18 weeks of age. I. Time spent in goal quadrant during Barnes maze probe test day. One-way ANOVA with Tukey’s post hoc. N=32 WT*^2Q/2Q^*, N=15 *f-ATXN1^146Q/2Q^*, N=19 *ATXN1_mKO_* mice 18 weeks of age. * p < 0.05, ** p < 0.001, **** p < 0.0001. Data is average ± SEM with individual mice presented as dots. Open dots represent female mice while filled dots represent male mice.

To determine how microglial *mATXN1* expression impacts SCA1 cognitive and mood deficits, we used open field, Barnes maze and fear conditioning.

Open field was used to assess anxiety-like behavior measured as decreased time in the center zone. While *f-ATXN1^146Q/2Q^* mice spent significantly less time in the center zone compared to WT controls, *ATXN1_mKO_* mice were indistinguishable from WT mice (Figure 8C) indicating that microglial ATXN1 contributes to anxiety-like phenotype in SCA1 mice.

Barnes maze was used to assess spatial cognition and problem-solving skills. Mice undergo four training days, each day they are placed on Barnes maze apparatus with four quadrants one containing an escape hole. Mice are encouraged to find the escape hole due to a bright light in the middle of the maze. Their movement is tracked and latency to escape times are measured. During training days only *f-ATXN1^146Q/2Q^* mice had increased latency to find the escape hole compared to WT mice (Figure 8D). To investigate why *f-ATXN1^146Q/2Q^* mice are impaired in finding the escape hole, we utilized Barnes maze unbiased strategy analysis (BUNS) of mouse path during assay to assess strategy development from random search to serial and spatial strategies over the four training days. Strategies (spatial>serial>random) are given a value based on cognitive requirement to develop and implement that strategy and the average value of strategies is used to calculate cognitive score. *f-ATXN1^146Q/2Q^* mice have a significantly lower cognitive score on the last training day compared to WT controls indicating deficits in strategy development that likely contribute to their increased latency to find escape holes (Figure 8E). Intriguingly, on the last training day *ATXN1_mKO_* exhibited increased serial strategy adoption compared to *f-ATXN1^146Q/2Q^* mice (Figure 8G, *bottom*). Indicating improvement in cognitive score and latency to escape times is due to *ATXN1_mKO_* mice shift towards serial strategy.

On the probe day, *f-ATXN1^146Q/2Q^* mice spent significantly less time in the goal quadrant compared to WT indicating memory deficits. *ATXN1_mKO_* were not statistically different from WT controls indicating improved memory (Figure 8I).

Previous studies identified impaired behavior in fear conditioning in SCA1 mice. Both *f-ATXN1^146Q/2Q^* and *ATXN1_mKO_* mice were similarly impaired in the fear conditioning assay (Supplemental Figure 5A-C), indicating that mutant ATXN1 in microglia does not contribute to impaired fear conditioning behavior in SCA1 mice. These findings indicate that microglial *mATXN1* significantly contributes to motor, cognitive and anxiety deficits in SCA1 mice.

## Discussion

Microglial roles in neurodegenerative disease have become increasingly appreciated in the past decade. Our results add to this body of work by interrogating the cell-autonomous nature of microglial activation in SCA1. This work is possible due to the recent advent of the conditional *f-ATXN1^146Q/2Q^* mouse model combined with human SCA1 iPSC derived microglia.

Our results indicate that *mATXN1* expression is sufficient to cause pro-inflammatory microglial phenotype in a cell-autonomous manner. We show increased reactivity in human SCA1 microglia through increased phagocytosis and release of pro-inflammatory cytokines including MIF, IL-16, IL-8, CCL1, CCL2, MIP-1α, CXCL-1, and CXCL12 in basal, untreated condition. Upon stimulation with LPS, SCA1 microglia undergoes a more pronounced shift to a pro-inflammatory phenotype (M1), compared to control microglia, characterized by enhanced secretion of pro-inflammatory cytokines and chemokines, including IL-1β, IL-6, and TNF-α. These results indicate that at both basal level and with inflammatory activation, human SCA1 microglia have increased levels of classic inflammatory markers and release of chemokines such as CCL1, CCL2, and MIP-1α, which are known to attract other immune cells (Garcia-Reitboeck et al. 2018; Wood et al. 2022) and may induce phagocytosis.

Additionally, we found increases in mitochondrial gene and protein expression in human SCA1 microglia. This is intriguing as most previous studies described mitochondrial deficits in SCA1 mice (Stucki et al. 2016). It would be of interest for future studies to identify why mutant ATXN1 causes seemingly opposite changes in mitochondrial gene expression in SCA1 human microglia and mouse SCA1 cerebella. We also identified a decrease in STING1 protein levels in SCA1 human microglia. Previous studies uncovered the role of mitochondrial stress to activate the type I interferon pathway through cGAS-STING (West et al. 2015). For example, heterozygous knockout of *Tfam* which is important for mitochondrial DNA maintenance (Larsson et al. 1998) resulted in increased interferon gene expression in a cGAS and STING dependent manner (West et al. 2015). Our proteomics data indicates reduced STING1 and normal levels of TFAM compared to control, indicating that increased mitochondrial gene and protein expression could result in increased mitochondrial activity to serve the energy demands of activated microglia. Further investigation is required to understand how microglia *mATXN1* expression leads to increased mitochondria genes and protein expression concomitant with decreased STING.

Intriguingly, in cerebella of *f-ATXN1^146Q/2Q^* mice we detected an increase in expression of ISGs: *Ifi47*, *Ifi28l2a*, *Iigp1*, and *Ifit1*, and interferon regulating transcription factors, which are significantly reduced in *ATXN1_mKO_* mice. This result suggests that mATXN1 may activate type 1 interferon pathway in cerebella of *f-ATXN1^146Q/2Q^* mice. Prior work has uncovered the role of cGAS-STING pathway in neurotoxic microglia. Cyclic GMP-AMP synthase (cGAS) is a DNA sensor found in cytoplasm and is a defense mechanism against viral DNA and self-genomic and mitochondrial DNA (Sun et al. 2013). Once DNA binds to cGAS, cGAS produces cGAMP which binds and activates stimulators of interferon genes (STING). Activated STING activates both interferon regulatory factor 3 (IRF3) and nuclear factor kB (NF-kB), a key transcription factor in neuroinflammation, ultimately leading to antiviral immune response (Zhang et al. 2020). Recent work in Huntington’s disease, another CAG expansion repeat disorder, indicated activation of cGAS-STING pathway (Sharma et al. 2020) caused by an increase in double stranded breaks resulting from mutant Huntington interfering with DNA repair (Sun et al. 2024). Additionally, work in ataxia telangiectasia uncovered increased self-single- and double-stranded DNA led to innate immune activation via the STING pathway (Song et al. 2019; Ferro et al. 2019). These studies add to a body of work revealing that malfunctioning DNA repair is a common feature among progressive movement dysfunction diseases (Ross and Truant 2017). Our RNA sequencing results similarly point towards a STING activation in SCA1 mice. One possible explanation for why there is an increase in STING in SCA1 mice is due to inability to repair DNA damage in SCA1. Previous studies have shown that ATXN1 localized to sites of DNA damage, and *mATXN1* impeded these dynamics hinting at a possible role in DNA repair (Stuart et al. 2021). On the other hand, we found that expression of mutant ATXN1 [43-46Q] in IPSC derived microglia resulted in significant reduction of STING protein levels. The difference in how human ATXN1 impacts human and mouse microglia could be a result of either increased CAG repeats in mice needed to model disease and cause DNA damage, and/or age/relative maturity of microglia, which also increases type 1 interferon gene expression (Roy et al. 2024; Gulen et al. 2023) and/or differences between species.

Comparing our human and mouse transcriptomics we show a shared downregulation of genes involved in mitochondria, the immune system disease and extracellular matrix indicating that microglia ATXN1 contributes to these changes in SCA1. We found that deleting mutant ATXN1 in microglia did not ameliorate perturbed expression of genes related to extracellular region, synapses, and development in SCA1 mice cerebella (Figure 6F), indicating that these changes are primarily caused by mATXN1 expression in neurons and other glial cells.

Importantly we found that microglial ATXN1 contributes to motor, cognitive and mood deficits in SCA1 mice. Interestingly, while removal of microglial *mATXN1* ameliorates motor coordination at late stages, it doesn’t impact early or middle stages of disease (Figure 8B). Previous work uncovered that preventing Atxn1 nuclear localization prevents rotarod phenotype from worsening (Handler et al. 2023). However, in that study preventing nuclear localization begins to prevent worsening rotarod phenotypes earlier on. Our results indicate that microglial *mATXN1* likely plays a role in the late stage; this is likely mediated through reduced late-stage immune response leading to improvements in neuronal health (Figure 5,6).

We found that microglial *mATXN1* plays an important role in anxiety-like phenotypes in SCA1 mice. Previous work indicates the microglia regulate anxiety through two separate microglia populations (Van Deren et al. 2025). Furthermore, SCA1 patient studies reveal increased anxiety (Fancellu et al. 2013; Brusse et al. 2011). Our work is the first to show that microglia *mATXN1* expression is important for SCA1 anxiety. There is reason to believe that increased microglia activation plays a role in anxiety, as patients undergoing interferon therapy for Hepatitis C virus show increased anxiety (Bonaccorso et al. 2001). Therefore, it is likely that increased anxiety in *f-ATXN1^146Q/2Q^* mice is due to increased interferon stimulated genes.

We found that microglial *mATXN1* expression does not improve fear acquisition or recall. However, our findings reveal for the first time that SCA1 mice are impaired in fear acquisition and recall, as previous work has uncovered that there is no difference in fear acquisition in *Atxn1^154Q/2Q^* mice, but there were impairments in fear recall, while *ATXN1[82Q]* mice exhibited impairments in fear recall but only on the last shock during acquisition. Importantly both *Atxn1^154Q/2Q^* and *ATXN1[82Q]* mice do not freeze significantly more than WT during novel context (Asher et al. 2020), indicating a unique fear conditioning phenotype in mice expressing humanized m*ATXN1* ubiquitously.

Additionally, we found that removal of microglial *mATXN1* improves cognitive score and serial strategy development during spatial learning in Barnes maze (Figure 8). Hippocampal neurogenesis is important for spatial tasks (Frechou et al. 2024) and microglial inflammation impairs neurogenesis (Sato 2015). Therefore, it is possible that reduced neuroinflammation in *ATXN1_mKO_* mice contributes to improved cognitive score and serial strategy development. While future work is required to fully understand the mechanism behind improvements in Barnes maze with microglial *mATXN1* reduction, our results contribute to SCA1 by emphasizing the importance of microglia autonomous *mATXN1* expression in SCA1 cognitive deficits.

One major limitation of this study is that *Lyve1^CRE^* mice express cre recombinase in microglial progenitor cells, macrophages, immune cells and lymphatic endothelial cells (Pham et al. 2010; Galanternik et al. 2017). Importantly, *Atxn1* is ubiquitously expressed, and previous work has uncovered *Atxn1* homozygous knock-out leads to activation of B cells which contributes to autoimmune injury (Didonna et al. 2020). Additionally, in acute injury both microglial and peripheral immune STING is neurotoxic resulting in cortical tissue loss (Fritsch et al. 2024). Therefore, it is possible that peripheral *mATXN1* expression contributes to disease state. If removing *mATXN1* from the peripheral immune system is the reason for the effects seen here, that would be a groundbreaking insight that would change our understanding of SCA1. To circumvent this limitation future work could utilize split-cre system which is highly specific to microglia or macrophages. This system has 40 amino acid peptides from the n-terminal behind *Sall1* promoter for microglia and *Lyve1* promoter for macrophages. The enzymatic 283 amino acid long c-terminal peptide is behind the *Cx3cr1* promoter. Expression of both is required for cre activity and is highly specific for microglia or macrophages, respectively (Kim et al. 2021).

In conclusion, our study has uncovered that cell-autonomous microglial *mATXN1* is important for microglial phenotype in both human and mouse SCA1. Furthermore, we elucidated increased mitochondrial gene expression, hypertrophy, increased phagocytosis and secretion of pro–inflammatory cytokines in human iPSC derived SCA1 microglia. We also found late-stage increase in interferon mediated inflammatory pathways in SCA1 mouse cerebella, which were corrected with the reduction of microglial *mATXN1*. Additionally, we found that *mATXN1* contributes to ramification and soma size in SCA1 microglia. Finally, we uncovered that microglia *mATXN1* contributes to cerebellar pathogenesis, motor, anxiety and cognition phenotypes in SCA1 mice. This work also helps further indicate the differences between human and mouse models of SCA1 disease, which should be considered when developing therapeutic approaches.

## Declarations Ethics approval

The study was conducted in accordance with the National Institutes of Health’s Guide for the Care and Use of Laboratory Animals and approved by the University of Minnesota Institutional Animal Care and Use Committee (2407-42222A , date of approval: 09/11/2024).

## Consent for publication

N/A

## Data Availability Statement

Data are available upon request.

## Competing Interests

The authors declare that they have no competing interests.

## Funding

This research was funded by NIH, grant numbers R01NS109077 (to MC) and R35NS127248 (to HTO). Adem Selimovic is supported by 1F31NS143172-01 training grant.

## Author Contributions

Conceptualization, M.C., A.S, and G.T., methodology, M.C., formal analysis M.C., G.T., A.S., G.F., S.C.R., E.P., I.M.B., and Y.Z.; investigation, G.T., A.S, V.S., and K.N.A. writing—original draft preparation, A.S., G.T., and M.C.; writing—review and editing, M.C., G.T., A.S., Y.Z, M.K., Y.N. and H.O., visualization A.S., G.T, G.F., and M.C.; supervision, M.C.; project administration, M.C.; funding acquisition, M.C. All authors have read and agreed to the published version of the manuscript.

## Acknowledgements

We thank all the members of Orr and Cvetanovic Labs for their help and suggestions. We acknowledge Genomic, Proteomics and Behavioral cores at University of Minnesota for their support.

---

**Supplementary Figure 1.** A. Flow cytometry analysis with CD43 antibody showing progressive acquisition of hematopoietic progenitor stages (iPSC-HPC) (CD43+) in unaffected control and SCA1 marking commitment to the myeloid lineage at the end of stage 2. B. Flow cytometry with CD45 and CD11b antibodies was used to validate microglia maturation. Flow cytometry analysis showing progressive acquisition of microglial surface markers during maturation. Representative histograms depict expression of CD45 and CD11b in unaffected control and SCA1 microglia.

**Supplementary Figure 2.** Flow cytometry analysis of pHrodo bioparticles uptake in SCA1 and control microglia as a percentage of pHrodo bioparticles with green-labeled cells.

**Supplementary Figure 3.** Overlap in DEG and TF in SCA1 imicroglia with SCA1 iMNs and microglia from single nuclei RNA seq analysis of SCA1 patient cerebella.

**Supplementary Figure 4.** Reduced human *ATXN1* expression in *ATXN1_mKO_* mice. A. RT-qPCR of human *ATXN1* expression. Unpaired Student’s t-test. B. RT-qPCR validation of microglia enrichment process. *Aif1* is a microglia marker, *Ncam2* is a neuronal marker, *Slc1a3* is an astrocyte marker. One-way ANOVA with Tukey’s post-hoc test. ** P < 0.01, **** P < 0.0001.

**Supplementary Figure 5.** Impact of selective deletion of mutant *ATXN1* in microglia on fear conditioning. A. Percentage of freezing after each shock during fear conditioning training day. * Indicates difference between f-ATXN1146Q/2Q and WT2Q/2Q and ^#^ indicates the difference between *ATXN1_mKO_* and *WT^2Q/2Q^*. Two-way ANOVA with Tukey’s post hoc test. * or ^#^ P<0.05, ** or ^##^ P<0.01, *** or ^###^ P<0.001, **** or ^####^ P<0.0001 B. Duration of freezing during the fear context during test day. C. Duration of freezing during novel context during test day. One-way ANOVA with Tukey’s post-hoc test. N=30 *WT^2Q/2Q^*, N=13 *f-ATXN1^146Q/2Q^*, N=11 *ATXN1_mKO_* mice 18 weeks of age. * P<0.05, ** P<0.01, **** P<0.0001. Data is average ± SEM with individual mice presented as dots. Open dots represent female mice while filled dots represent male mice.

## Notes

### Competing Interest Statement

The authors have declared no competing interest.

### Summary of Updates

The figures by adding figure titles and adjusting placement

