## Supplementary Figures for "Mutant ATXN1 impacts human and mouse microglia and contributes to cognitive, mood, and motor deficits in SCA1 mice"

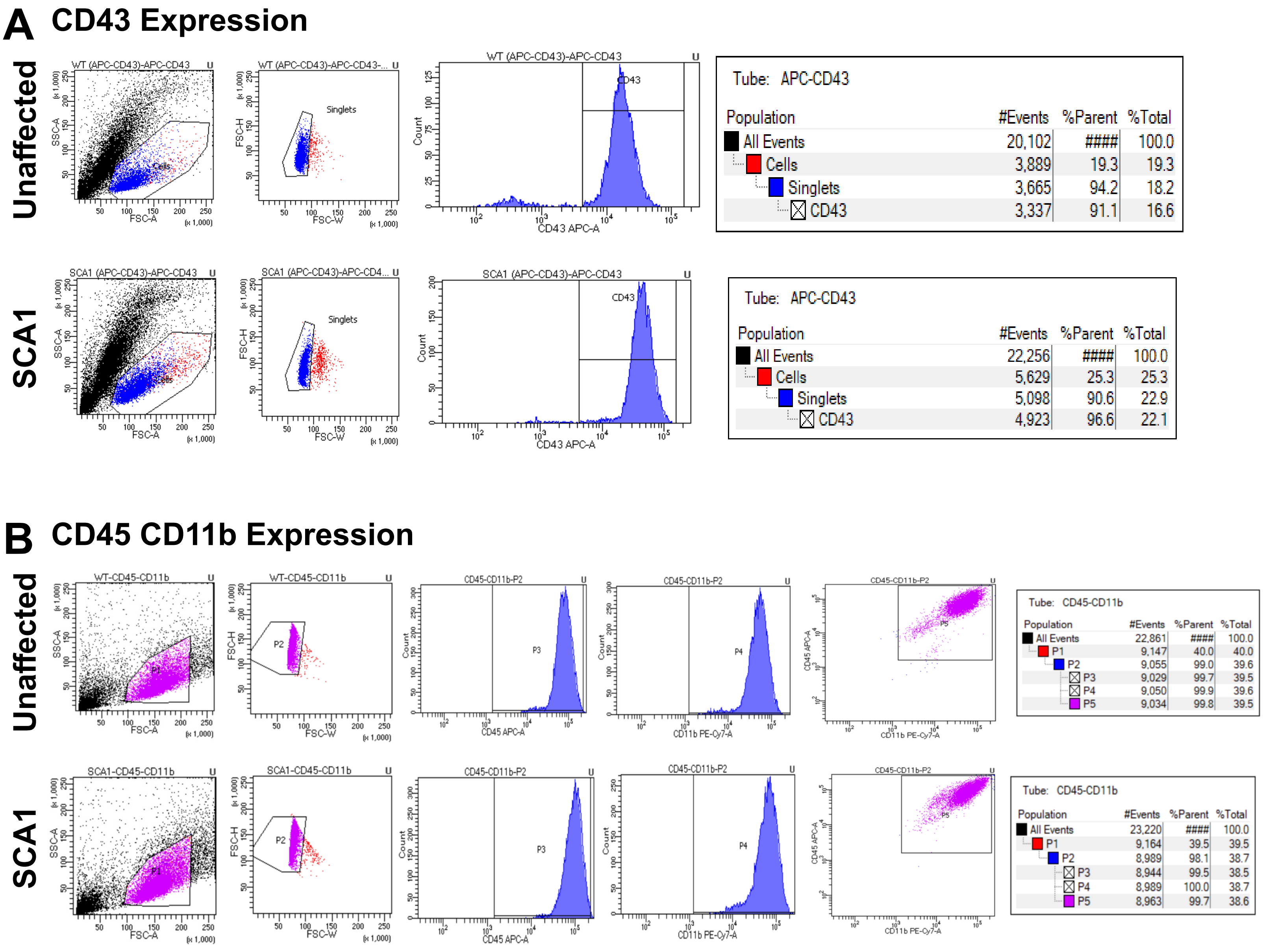


### Supplementary Figure 1

#### A unaffected control human microglia

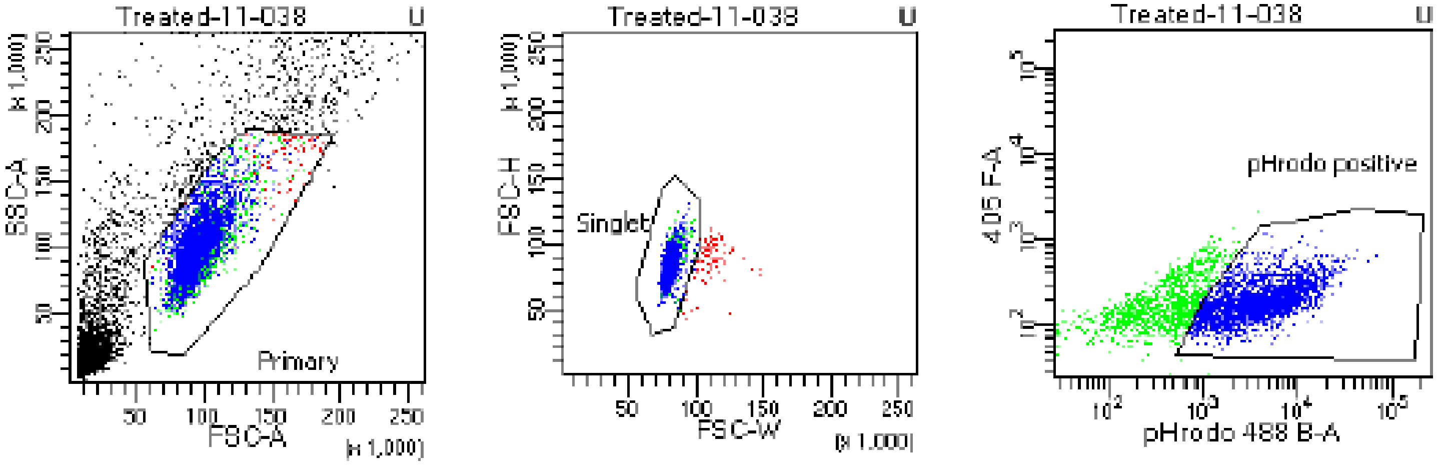


Tube: 11-038

| Population | #Events | %Parent | %Total |
|---|---|---|---|
| All Events | 10,000 | #### | 100.0 |
| Primary | 3,856 | 38.6 | 38.6 |
| Singlet | 3,768 | 97.7 | 37.7 |
| pHrodo positive | 2,610 | 69.3 | 26.1 |

#### B SCA1 human microglia

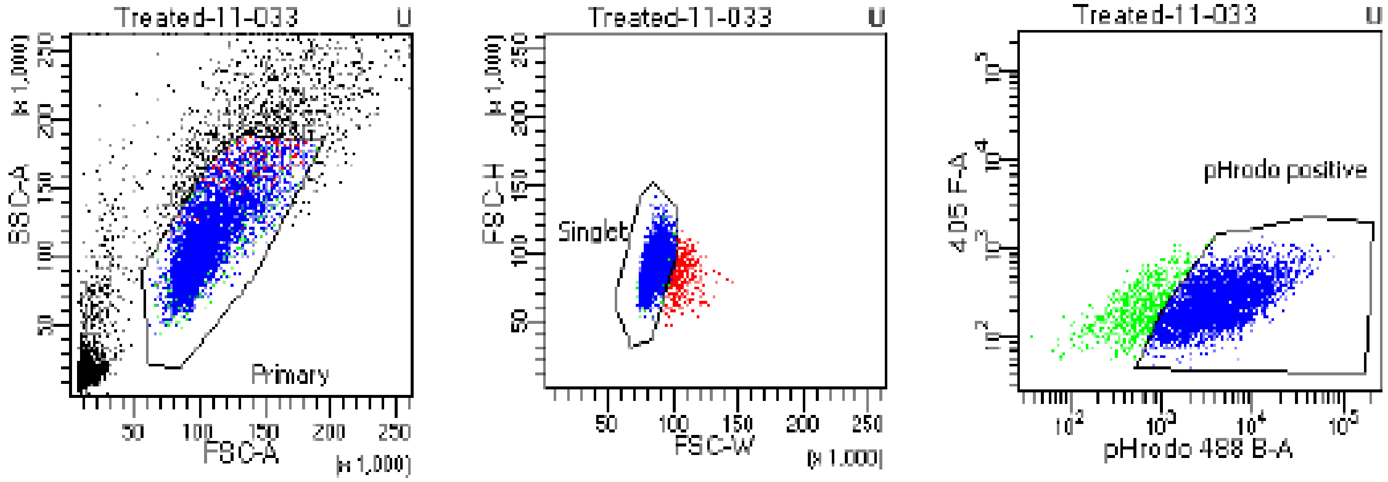


Tube: 11-033

| Population | #Events | %Parent | %Total |
|---|---|---|---|
| All Events | 10,000 | #### | 100.0 |
| Primary | 5,728 | 57.3 | 57.3 |
| Singlet | 5,374 | 93.8 | 53.7 |
| pHrodo positive | 4,651 | 86.5 | 46.5 |

### Supplementary Figure 2

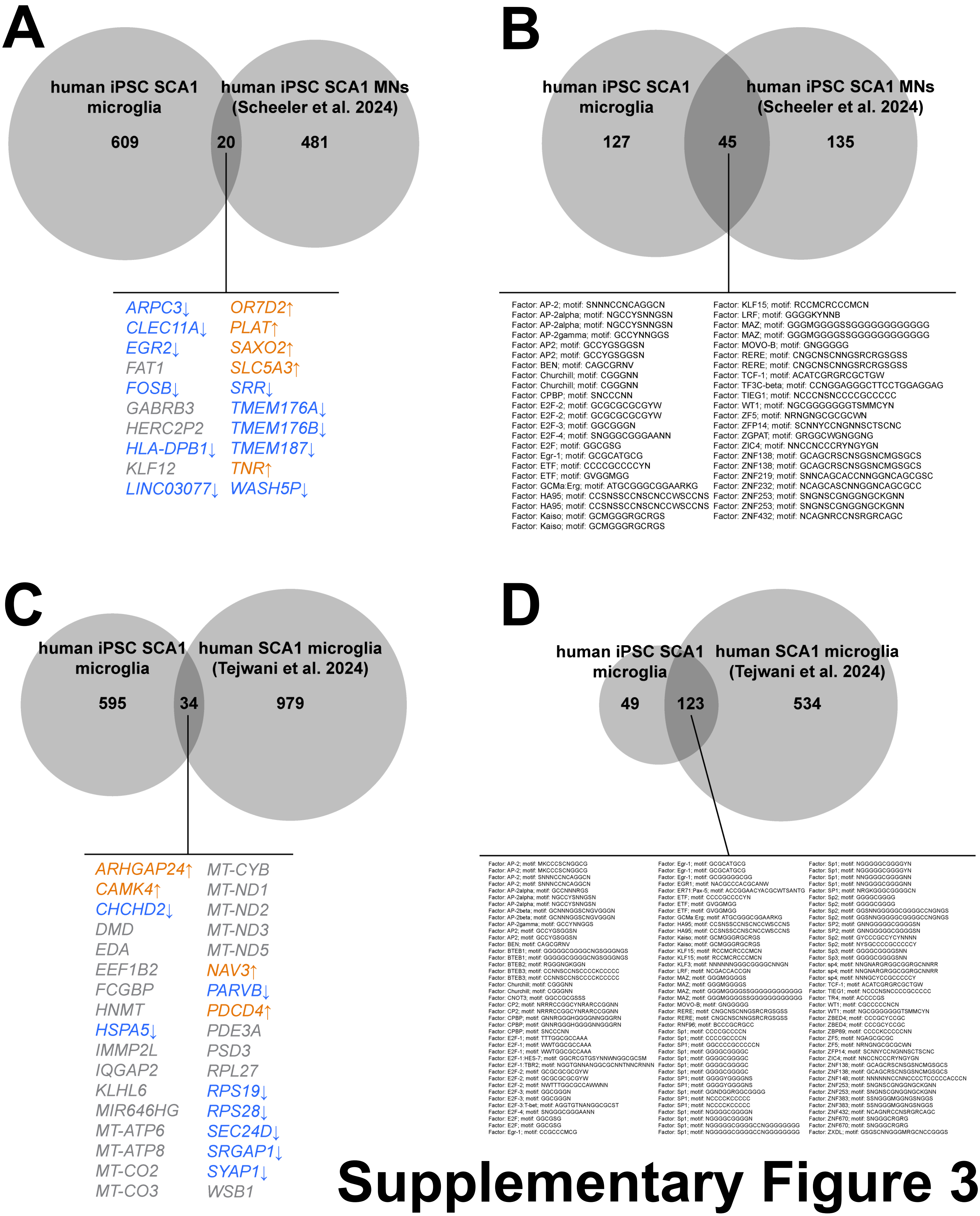


### Supplementary Figure 3

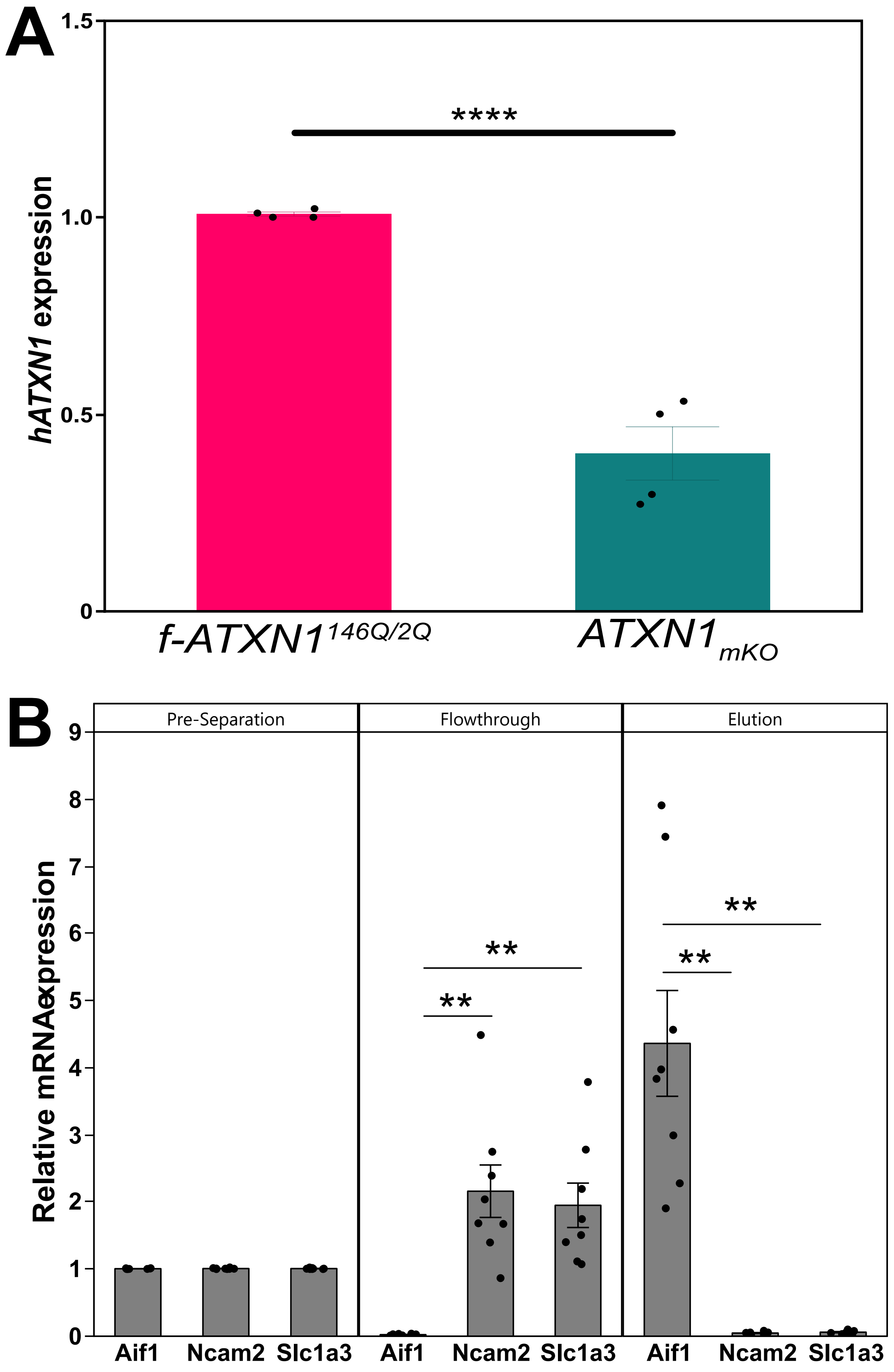


### Supplementary Figure 4

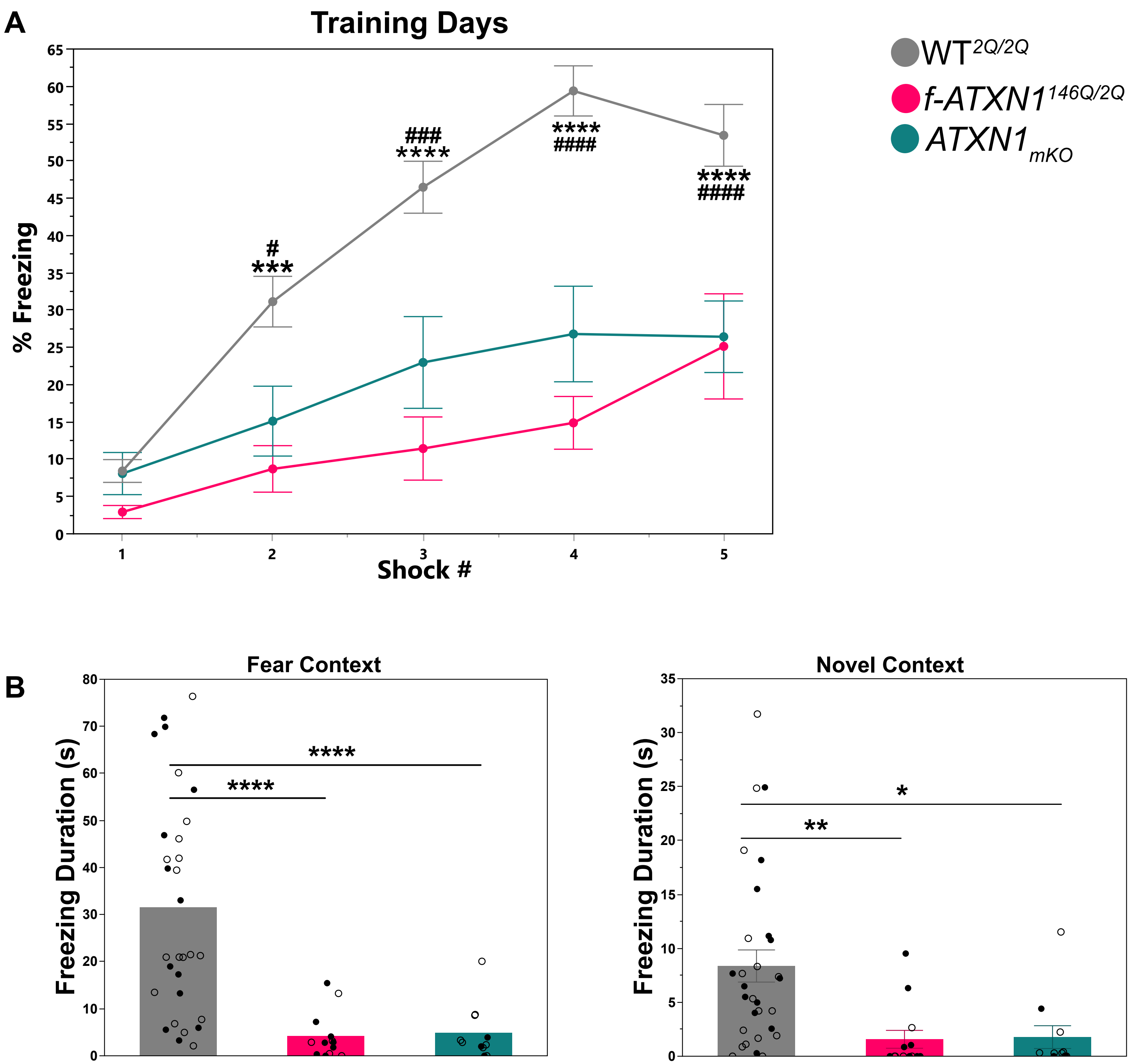


### Supplementary Figure 5
